# Systematic review and meta-analysis of *Hyalomma marginatum* vertebrate hosts in the EU

**DOI:** 10.1101/2024.09.11.612300

**Authors:** Madiou Thierno Bah, Laurence Vial, Lucas Busani, Léa Souq, Vladimir Grosbois, Celia Bernard, Ferran Jori

## Abstract

Host choice is a determining component of tick population and pathogen transmission dynamics. In Europe, ticks of the genus *Hyalomma*, which are involved in the transmission of several pathogens such as the Crimean Congo Haemorrhagic fever (CCHF) virus, are suspected to be spreading into new territories where they were previously unreported. Therefore, we performed a systematic review of the tick-host relationship of *Hyalomma spp* in Europe with a focus on *Hyalomma marginatum*, combined with a meta-analysis and meta-regression to describe its host preference pattern using three outcome values. Our initial qualitative analysis highlighted an increase in tick-host interaction rate in the last decades mostly in South-eastern and Central Europe. *H. marginatum* also appeared as the species holding the highest citation rate in terms of its association with hosts, and for which the largest number of host species were reported. The meta-analysis on *Hyalomma marginatum* host choice revealed preferential interactions for horses in the adult stage and birds of the *Emberizidae* and *Strigidae* families, in immature stages. Nevertheless, most of the heterogeneity of tick-host interactions remained unexplained suggesting the involvement of multiple drivers such as environmental or climatic conditions. Our results suggest that *H. marginatum* is a generalist tick whose distribution depends primarily on environmental conditions such as climate and habitat. Current limitations of our meta-analysis approach to identify hosts preference for *H. marginatum* and suggestions ways for improvement are further discussed.

## Introduction

Ticks are after mosquitoes, the second most important family of arthropod vectors involved in the transmission of vector-borne disease to humans. In Europe, the incidence of zoonotic tick-borne diseases such as anaplasmosis, Lyme borreliosis, Crimean-Congo Haemorrhagic Fever (CCHF) and rickettsiosis is increasing partly due to climate change (Gray et al., 2009).

Ticks (*Ixodidae*) are ectoparasites that require several hematophagous meals to complete their life cycle; thus, they alternate free-living and parasitic stages during their development. The non-parasitic stages include eggs, molting stages after blood-feeding, active or passive host searching stages after molting, and ovipositing females, are the most sensitive stages to environmental conditions. The transition from one development stage to the next one, except the emergence of larvae from eggs, requires blood feeding and therefore, the completion of a parasitic stage on a vertebrate host.

The range of host species on which a tick can feed varies depending on the development stages (e.g. usually immatures versus adults) and tick species. Most ticks require to feed on three-distinct hosts (to complete their development cycle (i.e. larvae, nymphs and adults feeding on different animals), but in some species the development cycle is completed on two (i.e ditropic) or even on a single (i.e monotropic) animal host. Some ticks are specialist while others are much more generalist. Historically, anatomical descriptions of tick mouthparts and coxae suggested that ticks had adapted to restricted groups of host species which they had co-evolved with, leading to the hypothesis that ticks were specialized parasites (Hoogstraal and Aeschlimann 1982). However, it was subsequently argued that constraints generated by environmental conditions during free-living stages were more critical to tick population fitness than those generated by host suitability and availability during parasitic stages, mainly because of the availability of a broad range of vertebrate host species to feed on (Klompen et al. 1996). This statement was later supported by studies showing that host community abundance and specific composition had a limited influence in the geographical distribution of tick species (Cumming 2002; Nava and Guglielmone 2013; Estrada-Peña et al. 2020). The most straightforward approach to assess whether a tick species is a specialist is to consider the number of host species it can feed on (Lymbery 1989). However, other criteria such as the prevalence of infested hosts, the intensity of infestation or the phylogenetic distance between parasite hosts have also been considered relevant (Poulin and Mouillot 2003; Rohde 1980).

The specificity or genericity in the host choice is important as it has a direct impact on pathogen transmission and the possibilities of disease spillover. Indeed, a generalist tick that could feed on a large diversity of hosts and whose geographic distribution is barely limited by host availability, can contribute to the emergence of diseases through geographic expansion or interspecific pathogen transmission (Mccoy, Léger, and Dietrich 2013; Dietrich et al. 2014). In addition, the density of generalist ticks is likely to increase as the number of species in the host community and/or the overall abundance of hosts increases. Because specialist ticks rely on a restricted number of host species and have a more limited potential in terms of geographic expansion, they are less likely to be involved in emergence events resulting from interspecific pathogen transmission or from tick colonization of new areas (Mccoy, Léger, and Dietrich 2013; Dietrich et al. 2014). However, because of their high level of adaptation to specific host species, populations of specialist ticks can locally thrive and contribute to the spread of their specific pathogens for which they are highly competent (Richard S. Ostfeld and Keesing 2000a).

In epidemiological systems involving a generalist vector, a vector-borne pathogen and multiple host species, the pathogen amplification transmission and maintenance potential may vary among the species of the available host community. In such systems, high species diversity can create a dilution effect resulting from the transmission of the pathogen to a wide range of hosts with low amplification, transmission and maintenance potential (Richard S. Ostfeld and Keesing 2000b; Schmidt and Ostfeld 2001). However, even if some hosts are poorly competent for the pathogen, they can still be good amplifiers of the tick and therefore contribute to boosting the tick population and thus, the potential number of infectious individuals. Therefore, there a less chances to observe a dilution effect in systems where the tick vector is a specialist species (Occhibove et al., 2022).

Tick species belonging to the *Hyalomma* genus are involved in the transmission of several important pathogens such as the Crimean Congo Haemorrhagic Fever (CCHF) virus. Immature *Hyalomma* specimens feed mainly on birds and small mammal species belonging the order *Rodentia* and *Lagomorpha* while the adults feed mostly on species belonging to the *Bovidae* family (Spengler and Estrada-Peña, 2018). However, the range and number of hosts involved in the *Hyalomma* spp. life cycle varies greatly across species. For example, *H. aegyptium* is a three-hosts tick that feeds mainly on *Testudo* tortoises (Široký et al., 2006). However, human and others vertebrates can act as occasional hosts. *H. dromedarii* is a two-hosts tick (occasionally one or three) mainly found on camels. *H. scupense* is a one or two hosts species that feeds preferentially on domestic ungulates, especially horses (*Equus ferus caballus*), and cattle (*Bos taurus*), regardless of its stage (Apanaskevich, 2004; Apanaskevich et al., 2013). Conversely, *H. rufipes* and *H. marginatum* tend to be more generalist (Estrada-Peña et al., 2018). Tick species belonging to the *Hyalomma* genus are involved in the transmission of several important pathogens of bacterial (rickettsial, anaplasma species), parasitic (theileria species) and viral origin such as the Crimean Congo Haemorrhagic Fever (CCHF) virus (Bakheit et al., 2012; Kumar et al., 2020). *Hyalomma marginatum* is one of the main vectors of the virus which is, currently considered as an emerging disease in Europe, Africa, Asia and the Middle East.

*H. marginatum* is a ditropic tick. This implies that larvae and nymphs feed on the same host, which is usually a small vertebrate (a lagomorph, a bird, a hedgehog or a rodent), whereas the adults usually feed on large ungulates such as horses, cattle, sheep, goats, wild boars, cervids or occasionally humans (Apanaskevich 2004). Considering its wide host range and its wide geographical distribution, *H. marginatum* could be considered as a generalist tick. However, host preferences have been reported at least for the adult stage (Grech-Angelini et al. 2016).

Furthermore, considering the hypothesis of “global generalist and local specialist” recently developed by MCcoy, Léger, and Dietrich (2013), ticks would locally develop preferences and adaptations to the most largely available host species leading to variations in the host species preference patterns across their distribution range. Under this hypothesis, an assessment of host preference at a wide scale would lead to falsely consider a tick species as generalist. Conversely, observation at the scale of the tick species distribution range of higher tick densities on certain host species, could result from the confounding effect of climate and habitat conditions rather than from genuine tick host preferences, and could thus lead to false conclusions regarding its classification as a generalist or specialist vector.

In this study, we conducted a systematic review of tick-host interactions of the *Hyalomma* species present in Europe in order to characterize the host community interacting with *Hyalomma* genus in Europe. Subsequently, we implemented a meta-analysis specifically on *H. marginatum* in order to describe its host species range, assess its host preference pattern and discuss its classification as a generalist or as a specialist tick species. Finally, based on those results, we discussed the potential influence on the most often reported host species on the tick life cycle and on the circulation of its vector borne pathogens such as CCHF.

## Materials and methods

We followed a systematic review approach, with the goal to extract any information related to *Hyalomma* spp tick-host interactions in Europe available in the literature from 1970 to 2021. The systematic review was conducted following as closely as possible PRISMA (Preferred Reporting Items for Systematic Reviews and meta-analysis) guidelines (Page et al., 2021).

### Publication selection

To achieve that goal, literature searches were carried out in the following databases: BIOSIS, CAB, Embase, MEDLINE, PubMed, Scopus and Web of Science. Publications were selected through specific keywords for Title (K1), abstract (K2) and within the core text (K3). The first list of keywords consisted of an enumeration of all *Hyalomma* species that can be found in Europe (Estrada-Peña et al., 2018) which were combined into query K1. The second list of keywords consisted of a list of species common names, families or clades that could act as potential hosts of *Hyalomma* ticks and were integrated into query K2. The third list of keywords consisted of an enumeration of all the countries within Europe which formed query K3 (Table 1).

**Table 1:**
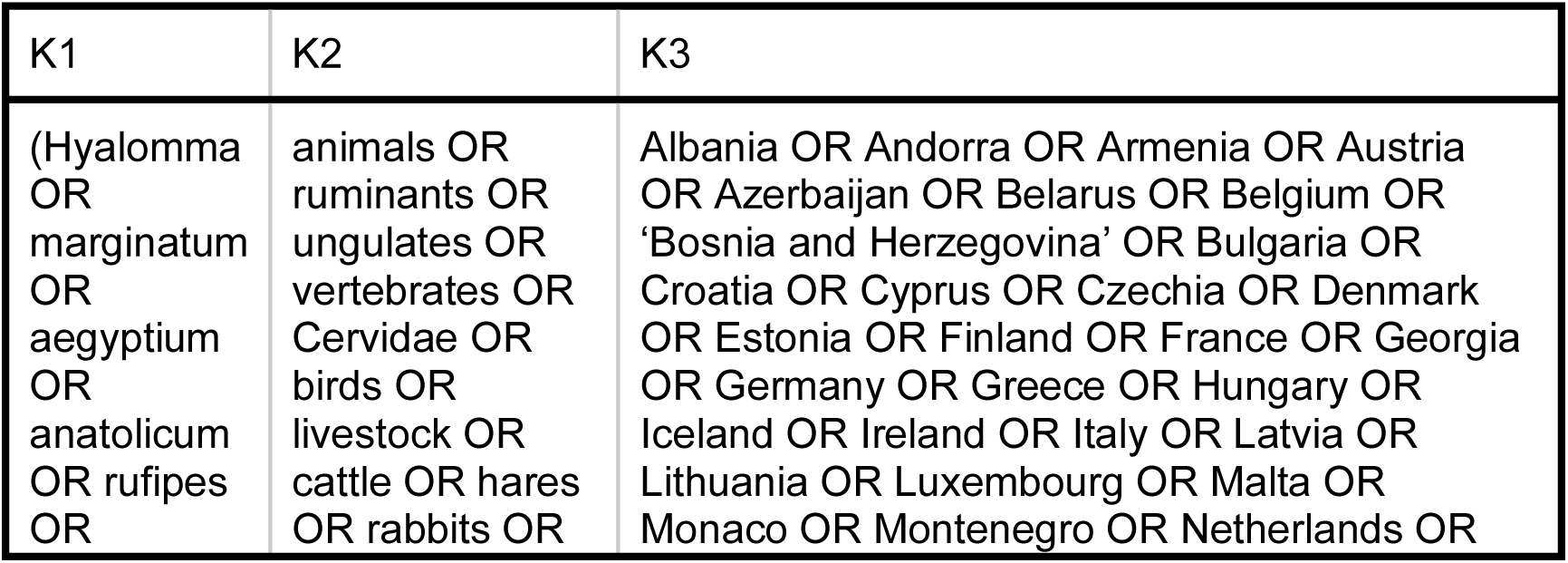
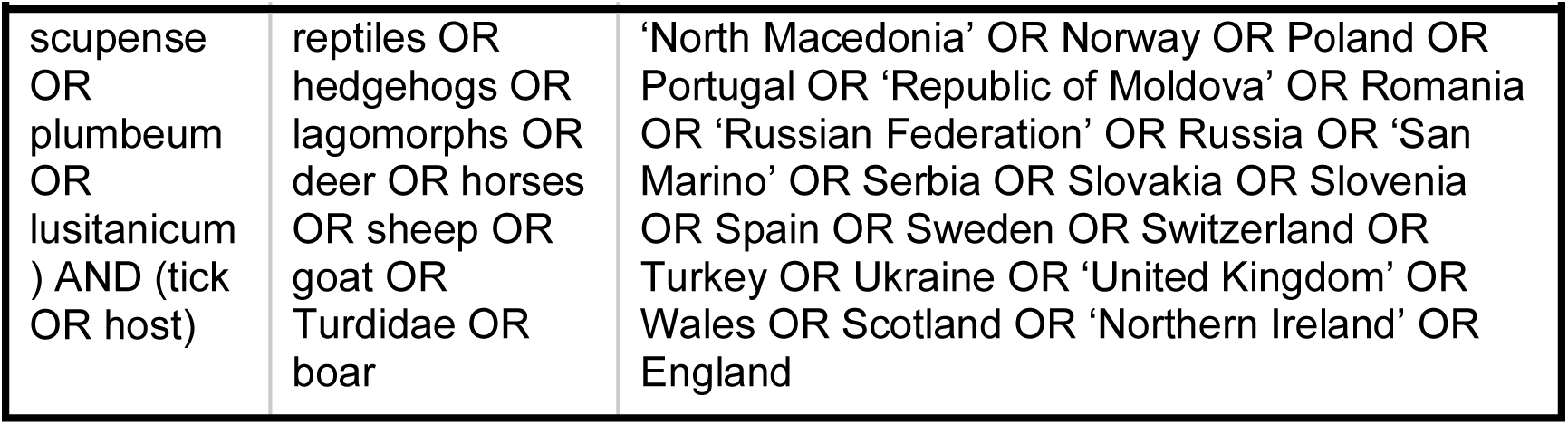
Queries used for the title (K1), abstract (K2) and body (K3) in the systematic literature review.

In addition to the keywords, inclusion and exclusion criteria were defined. Indeed, only original data collected in natural habitats on the trophic relationships of ticks of the genus *Hyalomma*, (i.e. a tick detected on a host), were included. As such, strict modeling studies, reviews and laboratory experiments were excluded. Moreover, we reduced our screening to publications which were in English or French. No exclusion criteria were applied related to the time of publication as it is supposed that host preferences should be constant across time, but we restricted the geographical area of the study to Europe. The choice of European countries was based on the list set up by the World Health Organisation’s Regional Office for Europe, which includes Europe in the broadest sense. In order to get a more homogenous geographical and environmental context, Central Asian countries (Kazakhstan, Kyrgyzstan, Tajikistan, Turkmenistan, and Uzbekistan) was considered too different from those in Europe considered, and therefore removed from the targeted list of countries chosen for the study. Finally, publications whose access was restricted (publications that were too old and present only in the archives, or whose full texts were not accessible) could not be included in our study.

Duplicates between databases were removed in Zotero (Roy Rosenzweig Center for History and New Media, 2022) and selected publications were imported into Rayyan, which is an open access software (Ouzzani et al., 2016) designed for the collaborative implementation of systematic reviews, scoping reviews and other knowledge synthesis projects, by speeding up the process of selecting studies through easy screening of titles and abstracts.

### Data extraction and compilation

After careful evaluation of the inclusion criteria in Rayyan, a full-text review was conducted on selected articles to extract pertinent data. A first database was compiled including the following qualitative data: title, authors, date of publication, country region or sub-region of the study and if available, date of sampling, host species and *Hyalomma* tick species involved, life stages of the sampled ticks as well as a classification of the invasive status of the collected ticks (imported or resident). The compilation of this shortlist allowed us to extract the following additional quantitative variables for each sampled host species: number of examined hosts, number of hosts infested by *H. marginatum*, number of *H. marginatum* ticks collected, and number of ticks collected in total. This data-extraction step was verified and validated by two different team members to reduce potential study miscount and misclassification errors.

In addition, to complete the geographical information related to each study, we extracted a measure of the most dominant landscape in which the sampling was conducted as well as the suitability of this habitat for the occurrence of *H. marginatum*. In order to do so, we relied on two raster maps, respectively: LANMAP 3 and the *H. marginatum* Suitability Map in Europe from Estrada-Peña and Wint (2016). LANMAP is a European landscape classification with four hierarchical levels, using digital data on climate, altitude, geological material and land cover as determinant factors (Mücher et al., 2010). It provides 350 landscape types, at the most detailed level, with 14000 mapping units with a minimum mapping unit of 11 km2. For the meta-analysis, we used a level of detail sufficient for distinction between climates.

The *H. marginatum* Suitability Map in Europe (Estrada-Peña and Wint 2016), used MODIS satellite data of daytime land surface temperature and the Normalized difference vegetation index at a 1km spatial resolution as predictive covariates for tick’s occurrence in a statistical regression model. Values from these two raster maps were extracted at the lowest geographical scale reported in each publication. Specifically, we computed the mean value of the suitability of *H. marginatum* within the region/subregion or country sampled and we extracted the most dominant landscape within the sampling zone being the region/subregion or the country.

### Data analysis

A Meta-analysis was then carried out on data extracted specifically for *H. marginatum*. The quantitative outcome, also called effect size, from various separate studies was pooled together in a statistical model to provide an estimate of the combined effect size and error. The effect sizes from several studies were expressed on the same scale, and each effect size corresponded to a particular outcome. Moreover, effect sizes were weighted in such a way that precise estimates (those with lower sampling error) had a higher influence on the pooled effect size than inaccurate ones. We investigated three effect sizes: 1. Proportion of hosts infested by *H. marginatum* among hosts examined for ticks, as a proxy of infestation rate; 2. Proportion of *H. marginatum* ticks among the total number of ticks collected on the hosts, as a proxy for competitiveness of the tick species; 3. Mean number of *H. marginatum* ticks collected per host, as a proxy of the parasitic load.

We used a random-effects model to partially pool the effect size, given the diversity of contexts involved in the original studies. Indeed, even after accounting for sampling error, an additional source of variation between studies was expected (Senior et al., 2016). This variation is referred as heterogeneity. Some studies in our meta-analysis contributed more than one observed effect size as they could report interactions of *H. marginatum* with more than one host species. Thus, heterogeneity was measured among records of interactions of *H. marginatum* with a host species in a study (publication) rather than among studies per se. The heterogeneity variance τ^2^ was calculated using the restricted maximum likelihood estimator for continuous effect sizes and the maximum likelihood estimator for proportions (Viechtbauer 2005). Among the various measures of heterogeneity that can be quantified based on τ^2^, we used the I^2^ statistic. I^2^ can be conceptualized as the proportion of variation in effect sizes that cannot be assigned to sampling error (Higgins and Thompson 2002). Thanks to its benchmarks it makes comparison of heterogeneity between meta-analyses easy. These benchmarks define small, medium, and high I^2^ as 25%, 50%, and 75%, respectively (Higgins et al. 2003).

Hosts, climate and environmental suitability were used as factors to explain the heterogeneity among effect size values retrieved from the selected studies. Thus, subgroup meta-analysis and meta-regression were conducted. In order to have a meaningful numbers of observations per category, hosts were grouped by family rather than species (e.g. *Caprinae* for sheep and goat, *Cervidae* to include red deer (*Cervus elaphus*), roe deer (*Capreolus capreolus*) and fallow deer (*Dama dama*), and bird families instead of bird species. Subgroups meta-analysis for the categorical variables (hosts and climate), and meta-regressions for the continuous variable (Suitability) were conducted. In a subgroup meta-analysis, it is hypothesized that effect sizes from different studies do not stem from the same general population but are rather divided in subgroups, each of which has a unique distinct overall effect. These subgroup effects are estimated, and a null-hypothesis significance test is performed assuming the absence of differences between subgroups. Cochran’s Q statistic (Cochran 1954) was employed to measure the differences between the estimated effect sizes and compared to its theoretical χ2 sampling distribution under the null hypothesis. Since effect-size estimates of subgroups based on fewer than 5 studies could lead to inaccurate estimations of τ^2^ (Borenstein et al., 2021). we assumed that subgroups shared a common estimate of the among-studies heterogeneity, in order to minimize this bias in τ^2^ estimation.

In a meta-regression, the outcome (response variable) is predicted according to the values of quantitative explanatory variables, similar to multiple regression. The outcome is the effect size, and the explanatory variables are quantitative characteristics documented in the different studies that can have an impact on effect size. We conducted this analysis on adult and immature stages separately, as their hosts belonged to different zoological families.

As a significant proportion of the interaction records concerned birds, we investigated if their behavior (feeding, migration) could explain the observed differences in the three outcomes used in the meta-analysis, respectively infestation rate, competitivity and parasitic load. Bird behavior was condensed in a single parameter according to the feeding behavior and the migratory status of each species, as such they were classified in three categories: - 1 Birds spending time on the ground and not migrating - 2 Birds spending time on the ground and migrating - 3 Birds not spending time and the ground and migrating.

### Assessement of the potential Impact of bird behaviour

In order to correlate this behavior parameter to the three outcomes, we run three generalized linear model in the *brm* package after specifying informative priors for the model parameters. The prior distribution for behavior was modelled as a normal distribution (0,2), as we expected small estimate due to our limited number of studies. We employed a zero inflated beta regression to model the relationship between infestation rate and behavior. To meet model assumptions, we substracted each value of infestation rate from 1 as it was initially an excess of 1.

Concerning the relationship between competitivity and behavior, we employed a beta regression to model.

For the relationship between parasitic load and behavior we employed a quasi-poisson distribution for modeling.

The Bayesian regression model was expressed as: Y ∼ X1 + ε, where Y the response variable is one of the three outcomes and X1 is the behavior as a predictor variable.

We mostly used default *brm* settings, including four chains, which were run for 2000 iterations and discarding the first 1000 as a burn-in period. We used default priors except for behavior estimate, for which we selected a narrower prior considering the numerous confounding factor among the studies from which the values were extracted. Specifically, we used normally distributed priors with a mean of one and a standard deviation of 2.

For each metric, response we calculated the proportion of the posterior distribution of the mean comparative behavior estimate (corresponding to the intercept against one of the level of the factor) above or below zero, equivalent to the probability of an increasing or decreasing mean outcome comparatively to bird behavior.

The average marginal effect of each factor level of bird behavior was also investigated. Furthermore, to visualize the posteriors for birds behavior we used a hypothetical dataset to generate predictions using 4000 draws.

Statistical analyses were conducted using the R statistical software version 4.3.2 (R Core Team, 2023) with the packages *meta* (Schwarzer 2022), *metafor* (Viechtbauer 2022) and *brm* (Bürkner 2021).

## Results

### Bibliometric outputs

The initial query of the seven target data bases yielded 1448 records, which reduced to 435 articles after removing duplicates. Among these, 319 references were selected for full reading during the first phase of selection by title and abstract, using Rayyan. A total of 100 references could not be exploited, either because the full text was unavailable or because they were written in a language other than French or English. The remaining 219 articles were read, and three of them were finally discarded because they were considered irrelevant. The remaining 219 publications were included in the qualitative analysis, 190 of them providing sufficient details to be included in the quantitative analysis (Figure 1).

**Figure 1.**
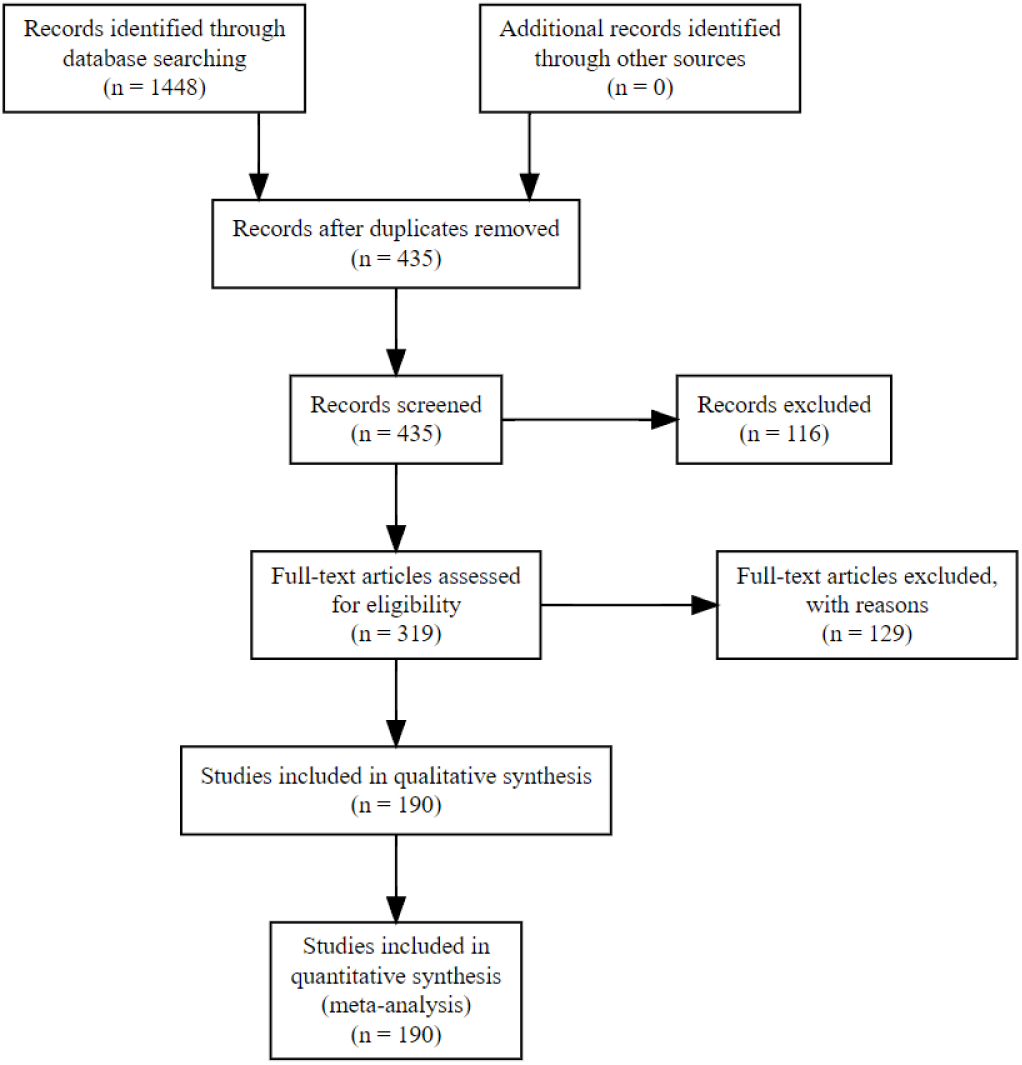
PRISMA flow chart representing the selection of studies included in the systematic review on the dispersal and maintenance of ticks of the genus Hyalomma in Europe by wild and domestic animals

Among the 190 publications, the oldest ones dated from 1969, and only thirteen were published between 1969 and 1999. Most of the selected articles (137) were published between 2011 and 2021 (Fig 2).

**Figure 2.**
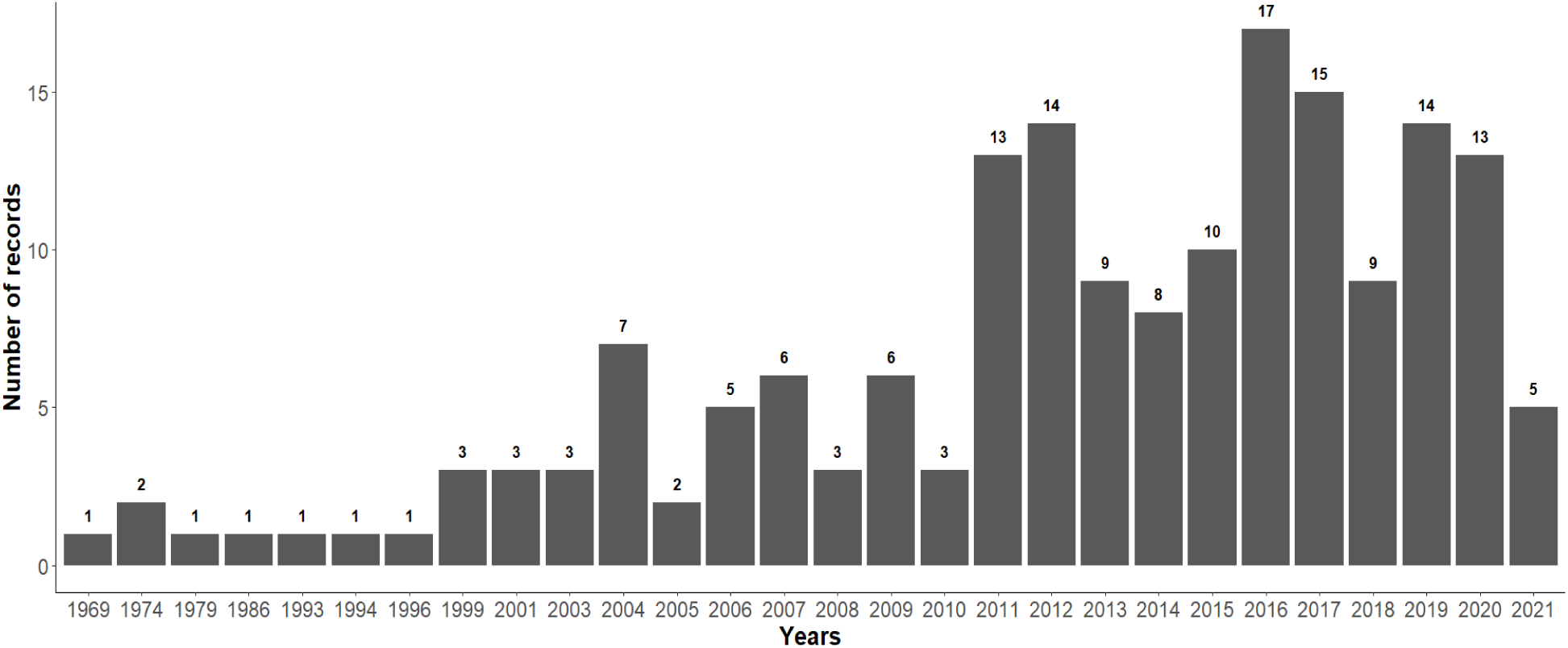
Number of publications selected for quantitative analysis per year included in the systematic review on the dispersal and maintenance of ticks of the genus Hyalomma in Europe by wild and domestic animals

The selected publications covered 29 countries. Among them, Turkey was the most represented with 23% of the selected publications, followed by Italy and Spain representing 16% of the selected publications each (Fig 3).

**Figure 3.**
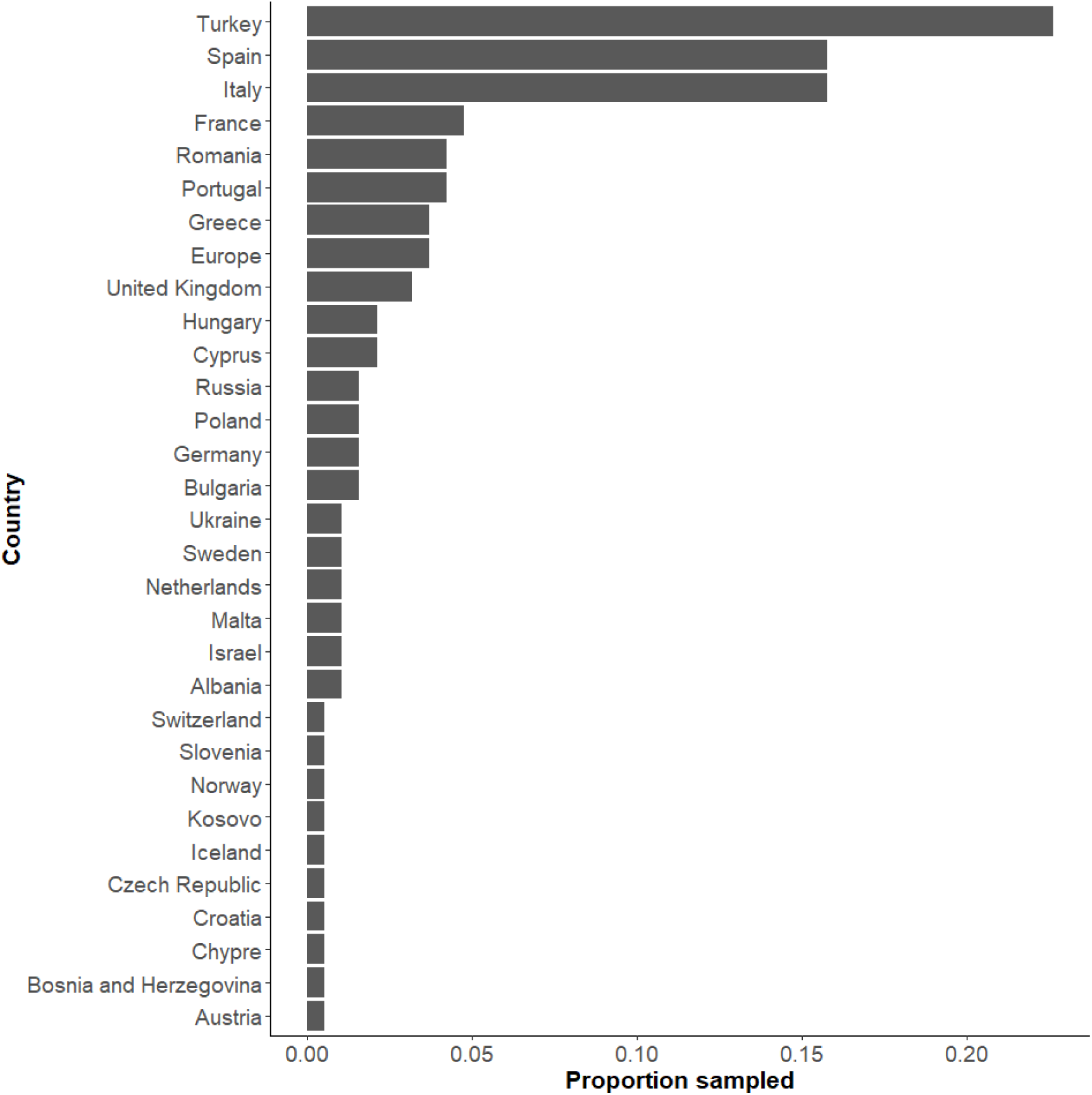
Proportion of publications selected for quantitative analysis by country

### Qualitative analysis on the different *Hyalomma* tick species

Among the 190 remaining publications, 85 papers (45%) reported only interactions with domestic hosts, 70 (37%) only with wild hosts and 24 (12%) with both (wild and domestic hosts). A total of 9 papers (5%) concerned ticks in free questing stages on vegetation and 2 (1%) were human case reports (Figure 4).

**Figure 4.**
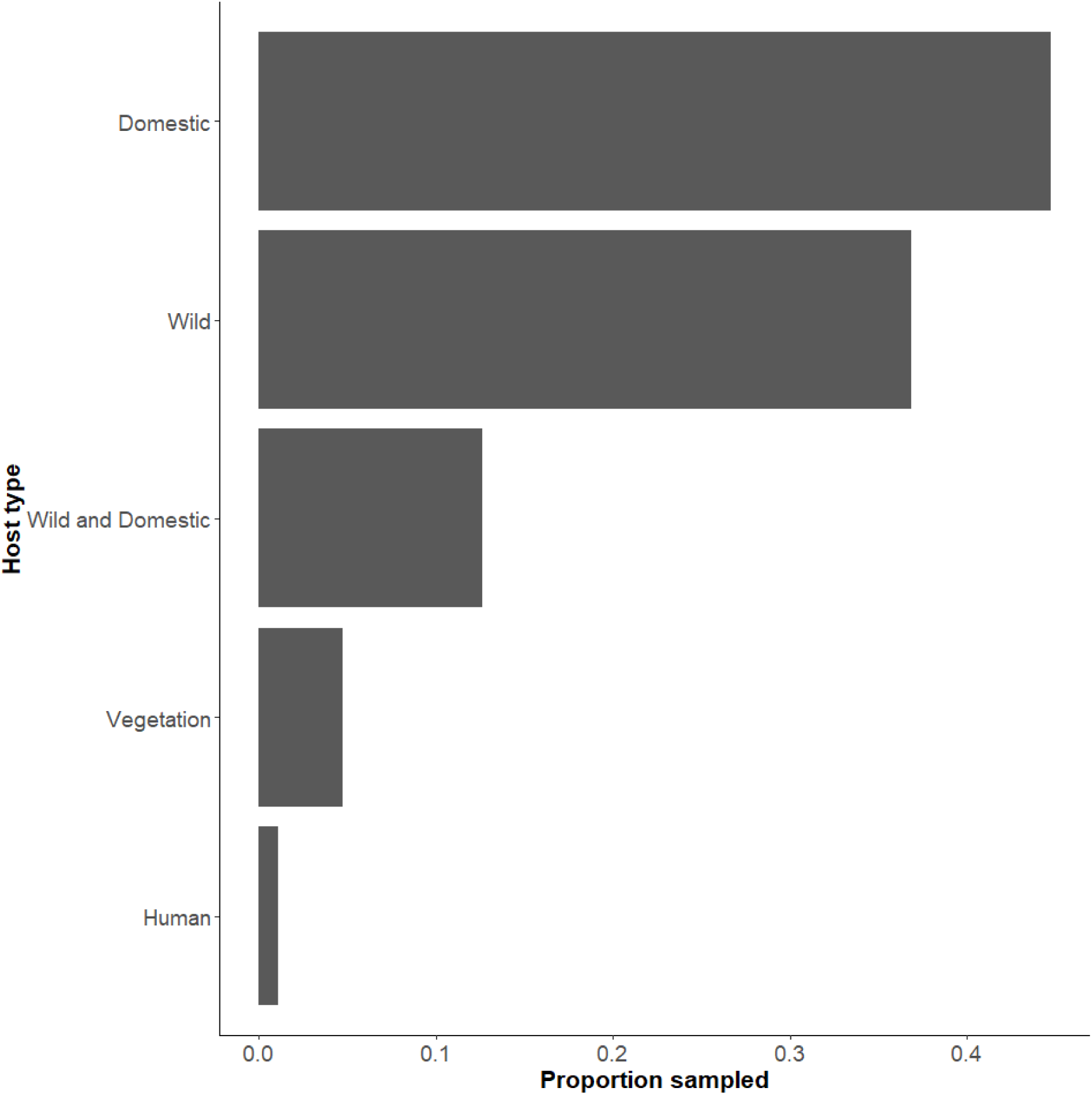
Sampling effort for each host type

Twelve species belonging to the *Hyalomma* genus were reported in Europe. The most common was *H. marginatum* with 142 citations (42%) (Table 2).

**Table 2:** Number of host species infestation distinct records for each *Hyalomma* species. Each citation represents one report of a tick species infesting one host species reported in a publication.

| Species | Number of records | Percentage |
| --- | --- | --- |
| *Hyalomma marginatum* | 142 | 41 |
| *Hyalomma excavatum* | 34 | 10 |
| *Hyalomma lusitanicum* | 32 | 9 |
| *Hyalomma rufipes* | 29 | 8 |
| *Hyalomma scupense* | 27 | 8 |
| *Hyalomma aegyptium* | 27 | 8 |
| *Hyalomma unspecified* | 22 | 6 |
| *Hyalomma anatolicum* | 18 | 5 |
| *Hyalomma dromedarii* | 3 | 1 |
| *Hyalomma asiaticum* | 2 | 1 |
| *Hyalomma impeltatum* | 2 | 1 |
| *Hyalomma truncatum* | 2 | 1 |
| *Hyalomma turanicum* | 2 | 1 |

Among *Hyalomma* species, the number of distinct reports of interactions with hosts and the number of reported hosts species were positively correlated. As such, *H. marginatum* was the *Hyalomma* species with the highest number of reported host species in our study (Figure 5). The second was *H. rufipes* even though its number of distinct records of interactions with hosts was relatively low (n=27). For the remaining five commonly reported *Hyalomma* species, the number of hosts was below twenty, possibly due to their lower number of distinct records of interactions with hosts.

**Figure 5.**
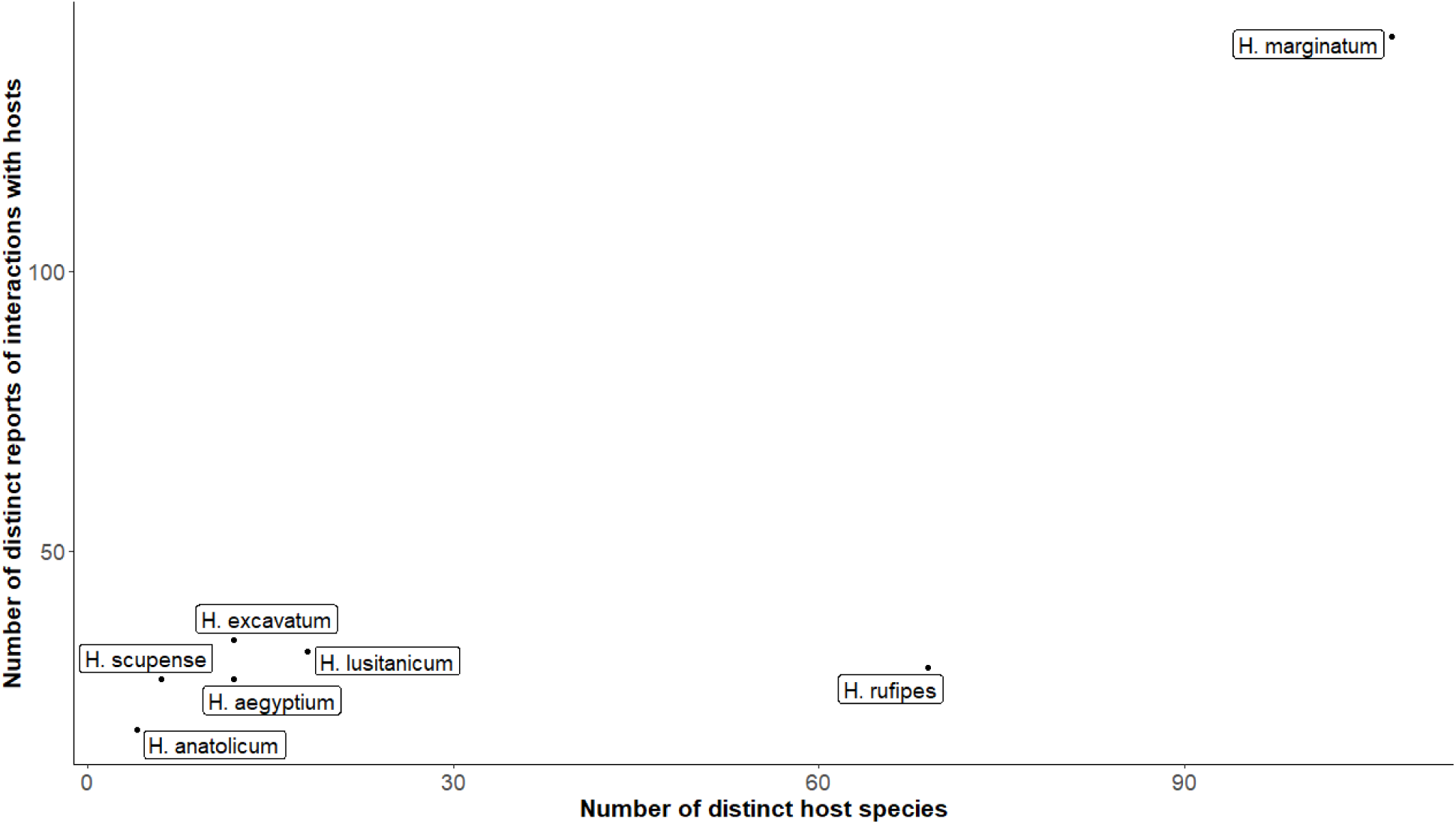
Number of distinct tick-host interactions reports in relation to the number of hosts distinct host species reported in Hyalomma tick species.

Figure 6 illustrates the distribution of *H. marginatum* across various host families, with the exception of the Agamidae family, indicating a broad diversity primarily driven by the presence of immature stages, particularly among avian families. *H. rufipes*, the second most frequently reported tick species, shows a narrower host range, primarily including mammal (*Equidae*, *Cervidae*, *Leporidae*, *Suidae*, *Erinaceidae*, *Canidae*, *Bovidae*) and bird (*Phasianidae*, *Turdidae)* families. On the other hand, *H. lusitanicum* was predominantly associated with wild host families such as *Cervidae* and *Suidae*, with limited reports within families representing domestic animals like *Bovidae* and *Equidae*. Similarly, *H. excavatum* was more associated with for wild host families, albeit with occasional reports on *Equidae*, *Bovidae*, and human hosts.

**Figure 6.**
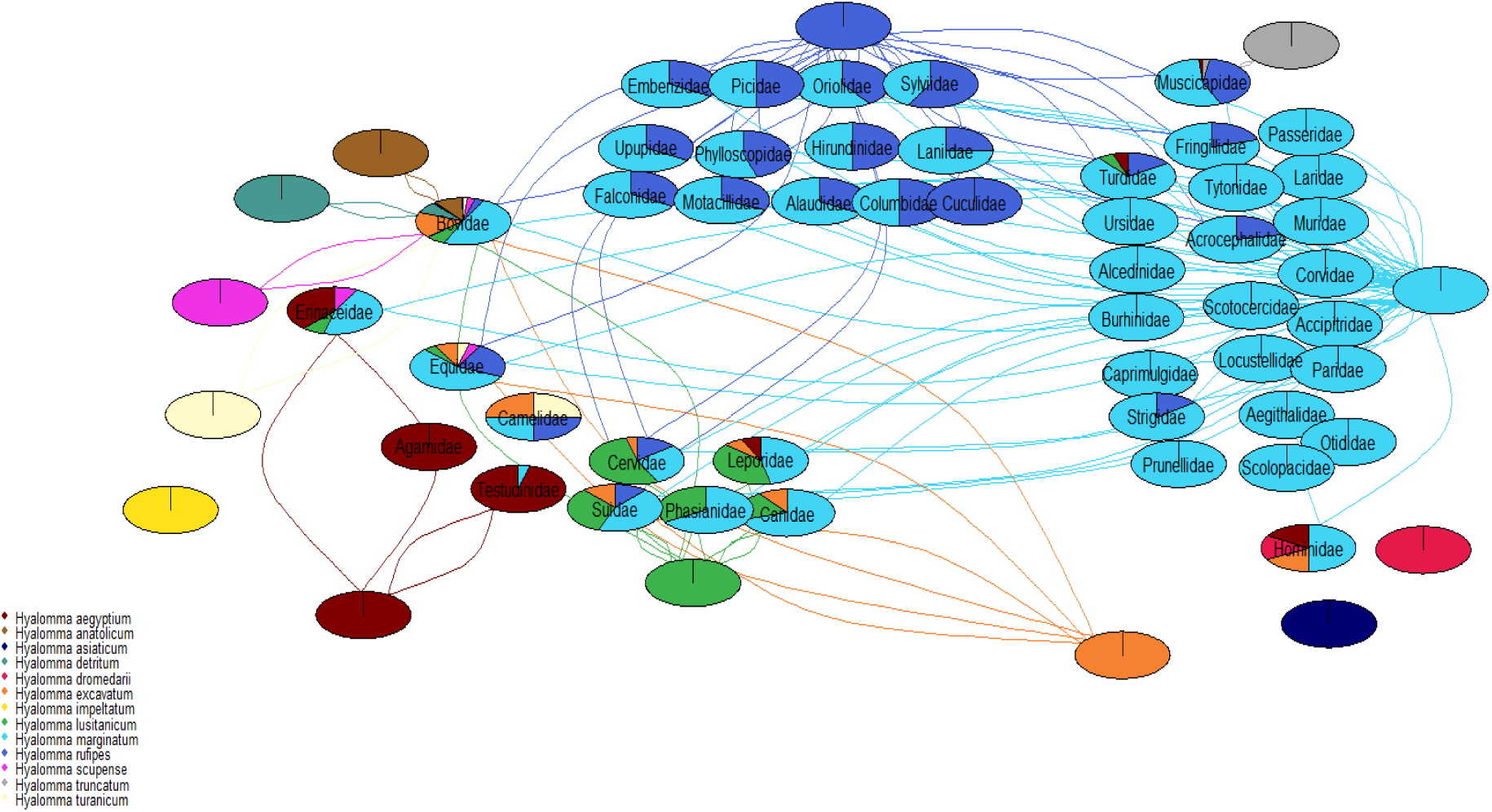
Network of tick-host interaction, in which each nodepie represents a family of hosts, and the proportion of each node pie is relative to the number of recorded interactions of the *Hyalomma* species of the same color. Only interaction which were reported at least two time are connected by an edge. *Hyalomma* species are not labelled as the legend describes the color associated with each *Hyaloma* species.

Fewer host families were reported for *H. aegyptium*, *H. anatolicum*, and *H. scupense*, as expected from the low number of distinct reports of interactions with hosts for these *Hyalomma* species. *H. aegyptium* was prominently associated with reptiles, particularly the *Agamidae* and *Testudinae* families, with additional reports in wild host families like *Leporidae* and *Testudinae*, along with sporadic reports on human hosts. Meanwhile, *H. anatolicum* and *H. scupense* primarily infest domestic hosts within the *Bovidae* and *Equidae* families, with *H. anatolicum* exclusively reported in the *Bovidae* family.

### Quantitative analysis on *H. marginatum*

Table 3 presents a summary of the effect sizes for the three predefined outcomes (infestation rate, competitiveness, and parasitic load) of the meta-analysis, estimated from the 75 relevant publications presenting quantitative data on the interactions between vertebrate hosts and *H. marginatum*. The number of reported interaction records differed greatly by development stage and outcome. On average, there were many more records for immature stages than for adult stages. The infestation rate was less reported than the two other outcomes for both stages. The estimates of effect size were quite similar across developmental stages, except for mean competitiveness, which was considerably higher in immature stages than in adult stages. Regardless of the effect size, confidence intervals are large and the heterogeneity between tick host interaction records was substantial (Table 3), which suggests the existence of several factors including host preferences or sampling biases, which deserve further investigation.

**Table 3:** Summary of the effect-size estimates with their 95% confidence interval and the estimated heterogeneity (I^2^) for the three target outcomes by developmental stage. The infestation rate is measured by the proportion of hosts infested by *H. marginatum* among hosts examined for ticks; the competitiveness by the proportion of *H. marginatum* in relation to the total number of ticks collected on the hosts; and the parasitic load by the average number of *H. marginatum* collected per host. Results are broken down by stage, as *H. marginatum* is predicted to feed on very different hosts in immature and adult stages, as all ditropic ticks.

| Development stage | Effect size | Number of interaction records | Estimated effect size | 95% Confidence Interval | Among interaction records heterogeneity ($I^2$ in %) |
| --- | --- | --- | --- | --- | --- |
| Adult | Infestation rate* | 14 | 0.106 | 0.003;0.296 | 90.7 |
|  | competitiveness** | 39 | 0.080 | 0.038;0.016 | 99.6 |
|  | Parasitic load*** | 36 | 1.625 | 0.59;2.661 | 99.8 |
| Immature | Infestation rate* | 45 | 0.091 | 0.057;0.141 | 80.5 |
|  | competitiveness** | 78 | 0.781 | 0.583;0.901 | 84.2 |
|  | Parasitic load*** | 63 | 0.064 | 0.045;0.083 | 78.3 |

\* proportion of animals \*\* proportion of ticks \*\*\* mean number of ticks per animal

### Host effect

#### Adult stage

We generally observed a relatively high imprecision (A max of 0.51 difference between the lower and upper estimates) in the estimated pooled effect sizes across tick-host interaction for each taxonomic regardless of the effect size (Figure 7). Infestation rate and competitiveness of *H. marginatum* differed significantly among host taxonomic groups (Respectively Q=98.44, p-value=<0.0001; Q = 138.30, p-value= <0.0001) (Supplementary S1; Supplementary S2). No significant difference was found among host taxonomic groups for parasitic load (Q= 5.15, p-value=0.2722).

**Figure 7.**
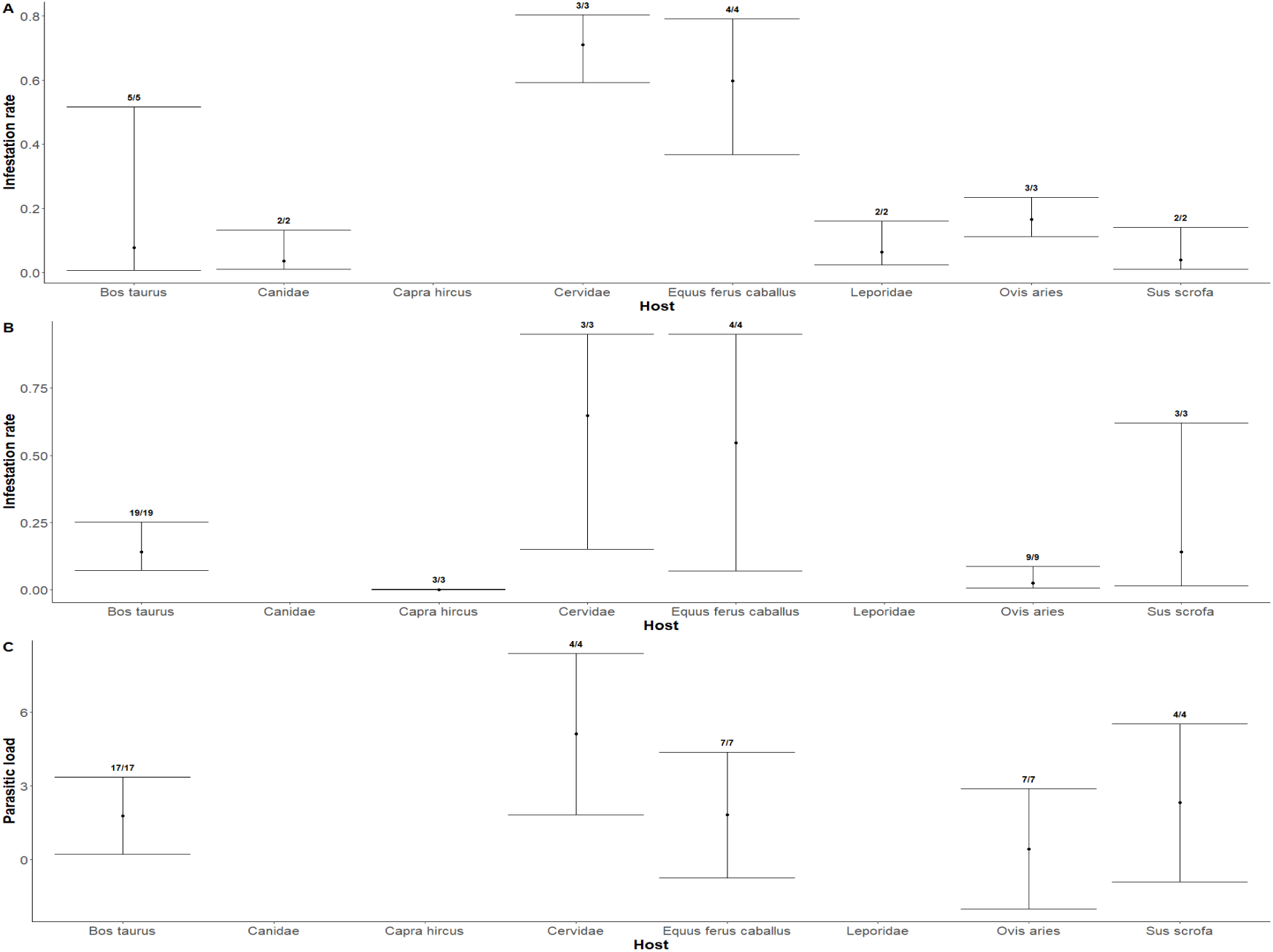
Effect of host taxonomic group on (A) infestation rate (B) competitiveness (C) parasitic load for adult stage. Effect size are represented with their 95% confidence intervals. The numbers on top of the error bar correspond to the number of distinct reports of interaction with host against the number of publications.

Competitiveness of *H. marginatum* was lower in *Caprinae* (*Ovis aries*, *Capra hircus*), *Canidae*, *Leporidae* and wild boar (*Sus scrofa*) in comparison to others host taxonomic groups.

Pooled effect sizes were generally higher in Horses, and Cervidae than in the other host taxonomic groups.

Residual heterogeneity I^2^ after accounting for host effect was 88.3%, 99.6%, 99.7% for infestation rate, competitiveness and parasitic load, respectively.

#### Immature stages

Similarly to the meta analysis for adult stages, we observed imprecise pooled effect size for most of the families of immature *H. marginatum* hosts regardless of the chosen effect size (Figure 8).

**Figure 8.**
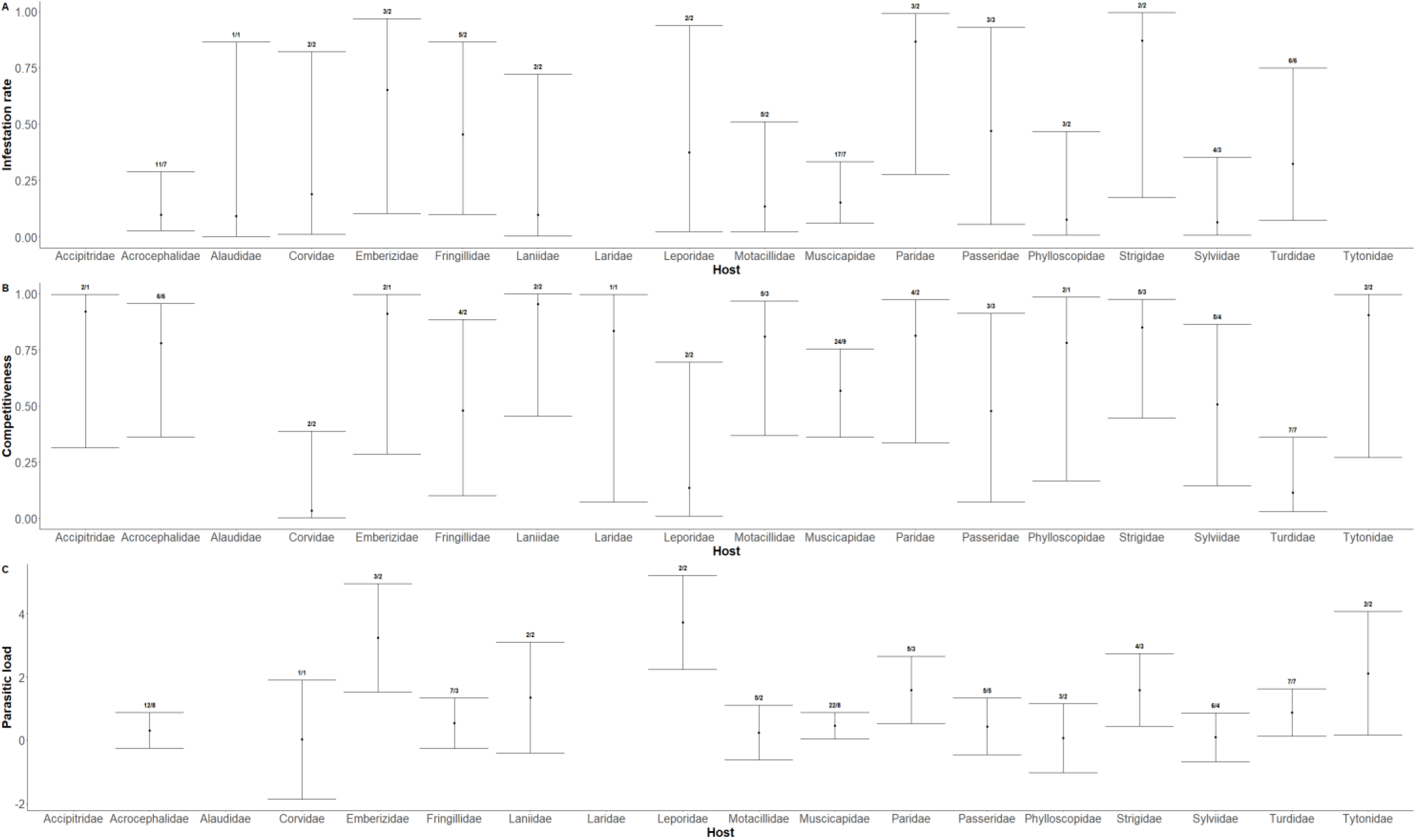
Effect of host species on (A) infestation rate (B) competitiveness (C) parasitic load for immature stages. Effect size are represented with their 95% confidence intervals. The numbers on top of the error bar correspond to the number of distinct reports of interaction with host against the number of publications

Significant differences were detected among host families for competitiveness and parasitic load (Respectively, Q=41.18, p-value=0.0003; Q=32.36, p-value=0.0090) (Supplementary S3; Supplementary S4)while no significant difference was found when comparing subgroups for infestation rates (Q=18.45, p-value=0.1871).

High parasitic load estimates were found for small mammals and birds such as *Emberizidae*, *Leporidae*, *Strigidae*, *Tytonidae* and *Paridae* families. High competitiveness estimates were found for most of the families except for the *Corvidae* and *Turdidae*.

Residual heterogeneity I^2^ after accounting for host effect was 83.1%,80%,86.2% for infestation rate, competitiveness and parasitic load, respectively.

### Landscape effect

#### Adult stage

The 95% confidence interval for pooled values for each effect size were wide (Figure 9). No significant difference was found between climates regardless of the effect size (Q=4.72, p-value=0.1935; Q=0.50, p-value = 0.9732; Q=1.19, p-value=0.9460 for infestation rate, competitiveness and parasitic load respectively). The highest estimation was found in Mediterranean and Anatolian climates, except for infestation rate where it was Continental and Mediterranean. Residual heterogeneity I^2^ after accounting for climate was 83.9%, 99.5%, 99.8% for infestation rate, competitiveness and parasitic load respectively.

**Figure 9.**
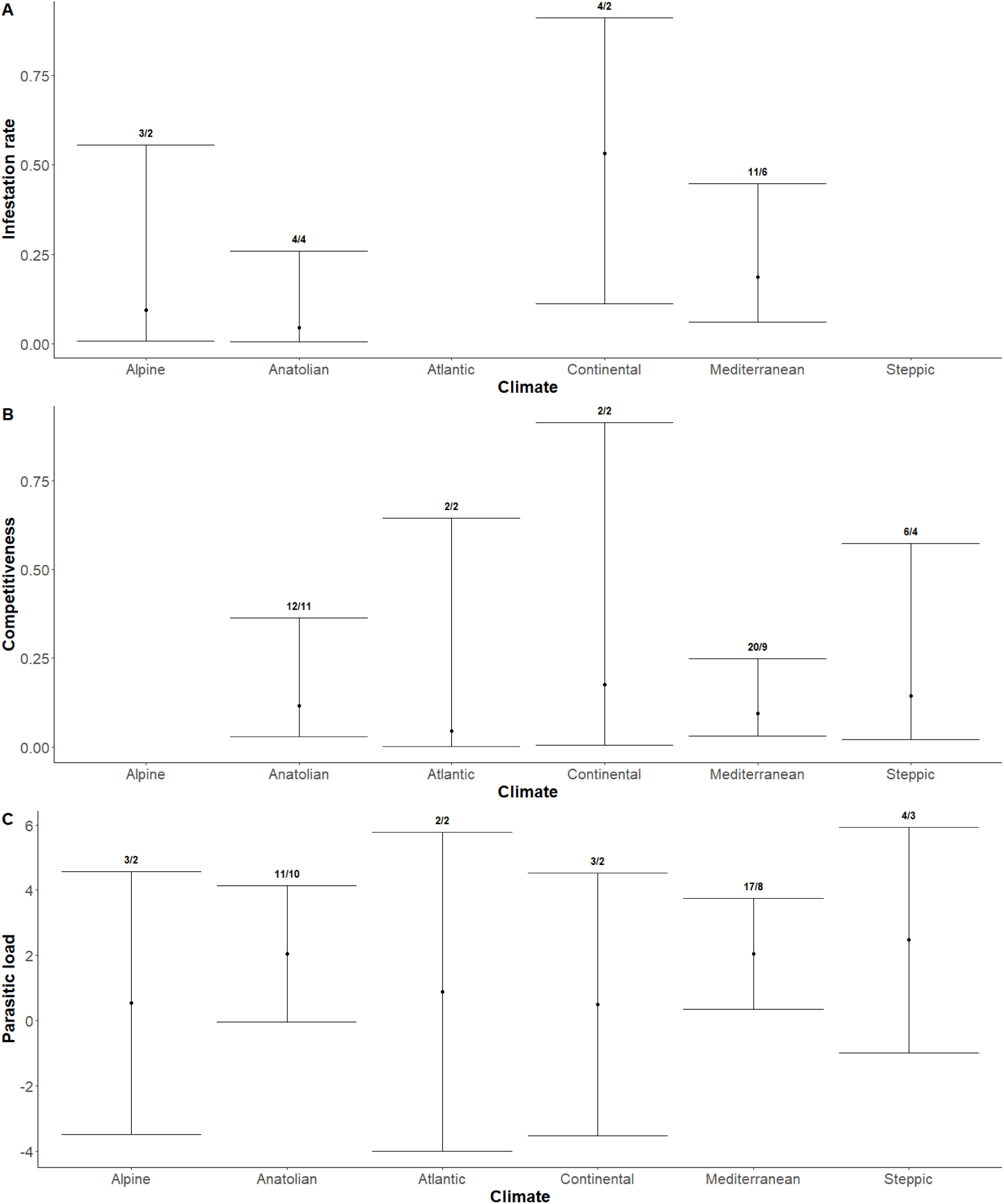
Effect of Climate type on (A) infestation rate (B) competitiveness (C) parasitic load for adult stages. Effect size are represented with their 95% confidence intervals. The numbers on top of the error bar correspond to the number of distinct reports of interaction with host against the number of publications

#### Immature stages

The 95% confidence interval were wide for pooled value of all effect sizes (Figure 10).

**Figure 10.**
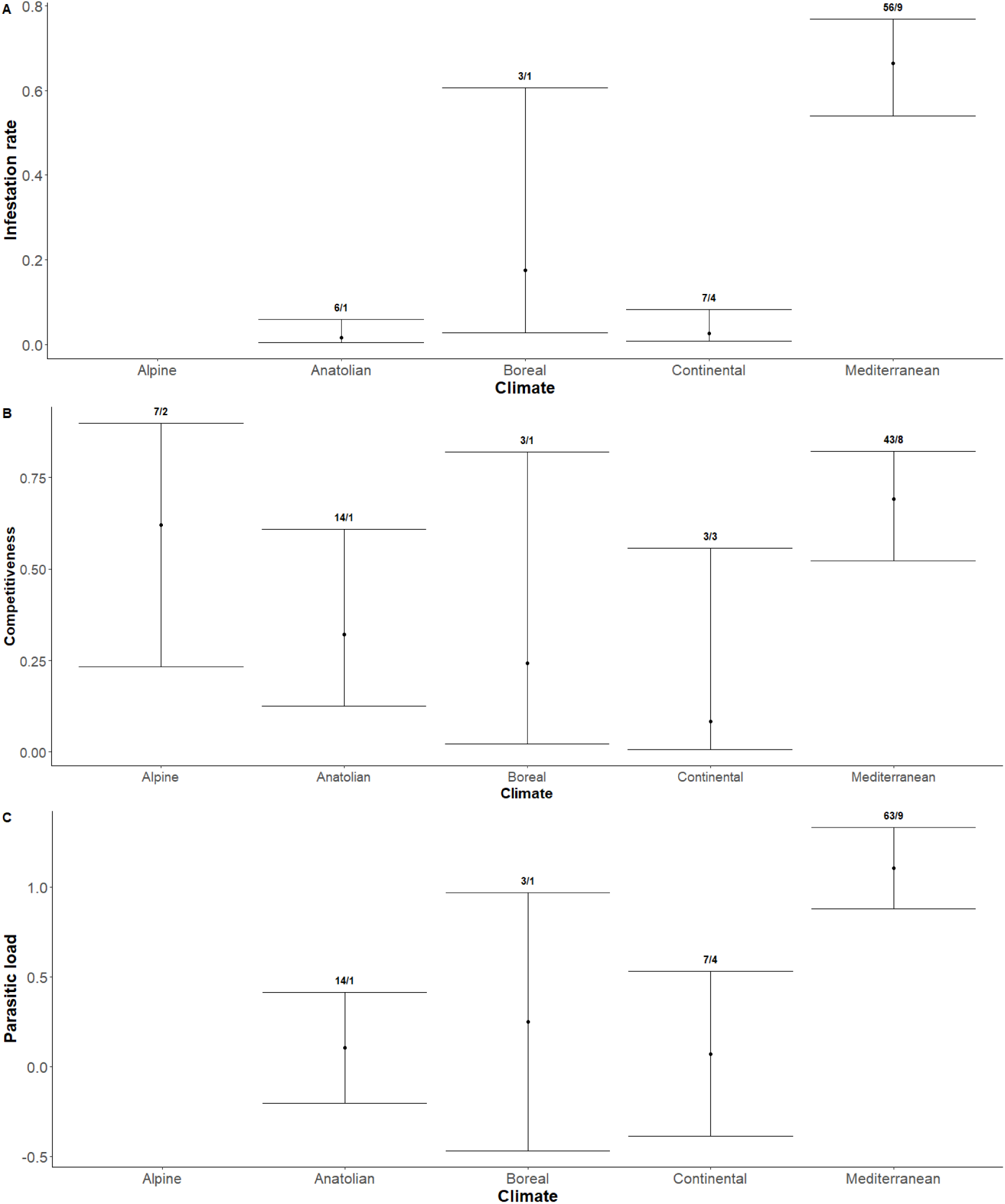
Effect of Climate type on (A) infestation rate (B) competitiveness (C) parasitic load for immature stages. Effect size are represented with their 95% confidence intervals. The numbers on top of the error bar correspond to the number of distinct reports of interaction with host against the number of publications

Significant differences were found among climates for infestation rate, competitiveness and parasitic load (Q=79, p-value<0.0001; Q=10.01, p-value=0.0403; Q=138.23, p-value<0.0001 respectively) (Supplementary S5; Supplementary S6; Supplementary S7).

Highest values were found in the Mediterranean climate for all effect sizes.

Residual heterogeneity I^2^ after accounting for climate was 87.4%, 92%, 71.0% for infestation rate, competitiveness and parasitic load respectively.

### Habitat Suitability effect

#### Adult stage

The residual heterogeneity I^2^ after adjusting for suitability was 94.8%, 99.90, 100% for infestation rate, competitiveness and parasitic load, respectively.

No significant effect of suitability was detected in adults ticks for infestation rate,competitiveness or parasitic load.

#### Immature stages

The residual heterogeneity I^2^ after adjusting for suitability was 95.26%, 91.49% and 99.99 % for infestation rate, competitiveness and parasitic load respectively.

No significant association was found for infestation rate and parasitic load Nevertheless, there was a significant positive association between habitat suitability and competitiveness during immature tick stages (QM = 13.9024, p-val< 0.0001).

### Effect of birds behavior on outcomes

Analysis of pairwise marginal effects across various bird behavior values, indicated that non-migratory species and those species feeding or hunting on the ground showed higher average levels of infestation rate, parasitic load, and competitiveness. Nonetheless, no substantial differences were observed between bird behavior classes (Table 4).

**Table 4:** Posterior distribution of the mean comparative behavior estimates. G/M correspond to ground feeding or hunting migratory birds; G/NM correspond to ground feeding or hunting non-migratory birds; NG/M correspond to non-ground feeding or hunting migratory birds.

| Contrast | Estimate | Conf.low | Conf.high |
| --- | --- | --- | --- |
| G/M - G/NM | 0.184 | -0.042 | 0.359 |
| NG/M - G/NM | 0.187 | -0.083 | 0.439 |
| G/M - G/NM | -0.254 | -2.24 | 0.756 |
| NG/M - G/NM | -0.773 | -2.69 | 0.332 |
| G/M - G/NM | 0.093 | -0.093 | 0.244 |
| NG/M - G/NM | 0.023 | -0.174 | 0.212 |

## Discussion

A better understanding of the interactions between ticks and their hosts is essential for population regulation processes and the spread of a tick-borne diseases. This study explored tick-hosts interactions within the genus *Hyalomma* with a focus on *H. marginatum*. Most of the publications retrieved from scientific publication databases were published between 2004 and 2021 suggesting an increased concern with regards to *Hyalomma* and associated diseases in Europe. A possible explanation for this trend could be related to climate change and modifications of land use that raise concerns regarding the expansion of *Hyalomma* ticks in Europe (Gray et al., 2009; Diuk-Wasser et al., 2021). Moreover, most of these publications referred to regions where CCHF has been documented to actively circulate such as Turkey, Mediterranean Islands, The Balkans region and Central Europe.

The high citation record observed for *H. marginatum* can be explained by the fact that it is the main vector of CCHF virus in many countries where the disease is endemic such as Turkey (Bente et al. 2013). Moreover, this tick is also the one for which the largest number of host species is reported. Nevertheless, it cannot be concluded that *H. marginatum* is the most generalist tick of its genus as this observation could be an effect of a more important sampling effort. Indeed, a correlation was observed between the number of distinct host-tick interactions records (which can be considered as a proxy of sampling effort) and the number of host species reported. This correlation has also been observed by other authors in the context of literature reviews to assess the link between specificity and sampling effort (Klompen et al., 1996). Noticeably, *H. rufipes* appeared as an outlier in this correlation, its number of citations being nearly 5 times lower than for *H. marginatum* but with almost the same number of host species reported (2/3 of the number for *H. marginatum*), suggesting its generalist character.

As expected, a strong heterogeneity among pooled effect sizes was found in the meta-analysis for all outcomes. Furthermore, as observed in previous systematic reviews, fewer infested species have been reported for adult stages than for immature stages (Nava and Guglielmone, 2013). This supports the hypothesis that immature stages are more generalist than adult ones due to their need to feed on morphologically larger hosts.

For adult tick stages, a significant effect of host species on the proportion of *H. marginatum* ticks infesting a host among all the ticks infesting that host (i.e. competitiveness) was detected. The effects of host species on the proportion of examined hosts infested by *H. marginatum* (i.e. the infestation rate) and on the average number of *H. marginatum* infesting the examined hosts (i.e. the parasitic load) were not significant. This can be explained by the limited number of exploitable records retrieved from the literature for parasitism effect sizes (resulting in wide 95% confidence intervals for the mean estimations of each host species). Nevertheless, higher estimates of these effect sizes were obtained for horses and *Cervidae* than for *Caprinae*, wild boars, hares and to a lesser extent, cattle. The estimations obtained for *Cervidae* must be considered with caution because the number of publications reporting values of effect sizes was low. In addition, in one of the publications, *H. marginatum* could have been confused with *H. lusitanicum* and misclassified. Aside from estimations for *Cervidae*, results obtained were consistent with one study where horses were more often and severely infected than cattle, wild boars and domestic goats from the same area in Corsica (Grech-Angelini et al. 2016).

Concerning immature tick stages, a significant variation among host families was detected for competitiveness and parasitic load with high mean estimations in the cases of rabbits and bird families (*Emberizidae Leporidae*, *Strigidae* and *Paridae)* for both effect sizes. *Emberizidae* encompass a small family of passerine birds which feed frequently on the ground, thus favoring contact with *H. marginatum*. On the other hand, high estimates of parasitism of this tick species were not expected for *Strigidae* and *Paridae* as species belonging to these families do not typically spend a large amount of time on the ground. However, estimations obtained for these two families were not particularly robust.

Despite record numbers and sample size issues, investigating variations in parasitism effect sizes at a zoological family level, may generate a certain within-group heterogeneity, since species belonging to the same zoological family may differ in their ecology and consequently in their availability and/or suitability as hosts for *H. marginatum*. Indeed, a strong intra-family heterogeneity translated into a large 95% confidence interval around the estimate, was observed with large differences in infestation rate or parasite load for certain species. For example, values of *Muscicapidae* family, ranged between 0.001 and 1.00 reported infestation rates values in the. This high heterogeneity can be partially attributed to interspecific differences within the family. We hypothesized that immature stages of *H. marginatum* behaved as generalist parasites of small vertebrates, implying that parasitism effect sizes would differ among potential host species, primarily due to differences in their availability influenced by behavior and habitat. However, our models did not reveal significant evidence supporting this hypothesis, despite finding higher *H. marginatum* parasitism estimates for non-migratory birds spending time on the ground, an activity that enhances host availability. This absence of effect could be explained by the limited availability of data for modeling and unaccounted confounding factors such as seasonality, geographical location, and sampling effort.

A consistent finding in our meta-analysis presented here is the large heterogeneity observed in effect sizes of parasitism that remain after adjusting for host species or host family effects. This suggests that other variables are more important in explaining *H. marginatum* variation in effect sizes than those considered in our study. In this regard, a substantial amount of heterogeneity was explained by the climate variable for immature stages with highest estimations obtained for the Mediterranean climate, supporting the results from previous studies on the drivers of the geographical distribution of *H. marginatum* (Bah et al 2022). Similarly, a positive relationship was found between immature *H. marginatum* tick competitiveness on hosts and environment suitability as derived from a statistical model of *H. marginatum* distribution fitted to *H. marginatum* presence-only data. These results support the hypothesis that the impact of climate might be more critical to the completion of *H. marginatum* immature stages than the composition of the potential host community (Cumming 2002; Estrada-Peña et al. 2020; Nava and Guglielmone 2013).

In contrast, in the analyses presented here, neither climate nor environmental suitability contributed to explaining heterogeneity in parasitism effect sizes for the adult stage except for the competitivity rate that had a positive correlation with the environmental suitability index.

The failure to detect significant relationships between parasitism effect sizes and environmental factors could arise from the failure to record accurate and precise information on geographical location, the dates of the surveys, the number of examined animals, the number of infested animals, and the number of ticks collected.

Another important limitation for identifying the relative influence of host species and environmental conditions, was the limited number of recorded data on the effect size of parasitism available in the identified sources literature for most of the potential host species or families. Given the large heterogeneity in effect-size values detected in most of the host species, which are likely to be related to environmental variability, mean estimates of effect size at species or family level were likely to be imprecise and biased (Borenstein et al. 2011). Moreover, effects of environmental conditions and host species or family are exposed to many potential confounding factors.

In order to better describe the tick-host relationship of *H. marginatum* using a meta-analysis approach, more abundant and accurate records of parasitism effect-size would be required for each potential host species or family. Moreover, in order to avoid confounding effects, such records should ideally originate from areas where environmental conditions are contrasted. In this particular context, surveys where ticks are recorded on several host species in the same area (ideally all the potential host species found in the area) at the same time of year are particularly useful for investigating the empirical variation of infestation levels among different hosts species. Using this type of approach would be much more efficient for addressing variation in infestation measures among host species than controlling for the potential confounding effect of the environment, since each record considered in the meta-analysis would reflect a difference in an infestation measure obtained from two different host species under the exact same environmental conditions.

In conclusion, our study highlights the relevance of assessing the abundance and diversity of the host community present in a given location in order to provide an estimation of the risk of pathogen transmission. From that perspective, our study suggests that *H. marginatum* is a generalist tick whose distribution depends primarily on environmental conditions such as climate and habitat. Its trophic relationships will therefore be primarily determined by the hosts community available in its immediate environment. The circulation of the pathogen will therefore depend on the number of hosts available and their capacity to replicate and amplify the virus and/or the tick population (Bernard et al. 2022; Occhibove et al. 2022). Moreover, our analysis highlights the importance of implementing as precise data collection and information on future surveys of tick species and their hosts that can be subsequently used for building statistical models to infer information on complex ecological host-vector pathogen interactions.

## Acknowledgements

We would like to extend our gratitude to William Wint for providing the suitability map for *Ixodes ricinus* and to Sander Mucher for supplying the LANMAP3 raster layers.

## Data, scripts, code, and supplementary information

The authors declare that they comply with the PCI rule of having no financial conflicts of interest in relation to the content of the article.

Data, scripts and code are available online: https://doi.org/10.5281/zenodo.13737782 Supplementary information is available online on biorxiv as part of the preprint.

