## Supplemental figures for "Systematic review and meta-analysis of *Hyalomma marginatum* vertebrate hosts in the EU"

### Supplementary

#### Host

##### Adults

| Study or Subgroup | Events | Total | Weight | IV, Random, 95% CI |
|---|---|---|---|---|
| **Hôtes = Bos taurus** | | | | |
| Grandi et al., 2020 | 1 | 1 | 3.7% | 1.00 [0.02; 1.00] |
| Aktas et al., 2004 | 1 | 2388 | 4.9% | 0.00 [0.00; 0.00] |
| Encinas Grandes, 1986 | 97 | 288 | 6.1% | 0.34 [0.28; 0.39] |
| Ica et al., 2007 | 61 | 300 | 6.1% | 0.20 [0.16; 0.25] |
| Koc et al., 2015 | 1 | 33 | 4.8% | 0.03 [0.00; 0.16] |
| **Total (95% CI)** | | **3010** | **25.5%** | **0.08 [0.01; 0.52]** |
| Heterogeneity: $Tau^2$ = 7.5895; $Chi^2$ = 66.4, df = 4 (P < 0.01); $I^2$ = 94% | | | | |
| **Hôtes = Canidae** | | | | |
| Encinas Grandes, 1986 | 1 | 25 | 4.8% | 0.04 [0.00; 0.20] |
| Grech-Angelini et al., 2020 | 1 | 31 | 4.8% | 0.03 [0.00; 0.17] |
| **Total (95% CI)** | | **56** | **9.7%** | **0.04 [0.01; 0.13]** |
| Heterogeneity: $Tau^2$ = 0; $Chi^2$ = 0.02, df = 1 (P = 0.88); $I^2$ = 0% | | | | |
| **Hôtes = Cervidae** | | | | |
| Chisu et al., 2017 | 1 | 2 | 4.1% | 0.50 [0.01; 0.99] |
| Keysary et al., 2011 | 4 | 4 | 3.9% | 1.00 [0.40; 1.00] |
| De La Fuente et al., 2004 | 46 | 65 | 6.0% | 0.71 [0.58; 0.81] |
| **Total (95% CI)** | | **71** | **14.0%** | **0.71 [0.59; 0.80]** |
| Heterogeneity: $Tau^2$ = 0; $Chi^2$ = 1.17, df = 2 (P = 0.56); $I^2$ = 0% | | | | |
| **Hôtes = Equus ferus caballus** | | | | |
| Jameson and Medlock, 2011 | 1 | 1 | 3.7% | 1.00 [0.02; 1.00] |
| Chisu et al., 2014 | 1 | 1 | 3.7% | 1.00 [0.02; 1.00] |
| Hansford et al., 2018 | 1 | 1 | 3.7% | 1.00 [0.02; 1.00] |
| Chitimia-Dobler et al., 2019 | 7 | 13 | 5.7% | 0.54 [0.25; 0.81] |
| **Total (95% CI)** | | **16** | **16.6%** | **0.60 [0.37; 0.79]** |
| Heterogeneity: $Tau^2$ = 0; $Chi^2$ = 0.74, df = 3 (P = 0.86); $I^2$ = 0% | | | | |
| **Hôtes = Leporidae** | | | | |
| Encinas Grandes, 1986 | 3 | 60 | 5.6% | 0.05 [0.01; 0.14] |
| Lorusso et al., 2011 | 1 | 7 | 4.7% | 0.14 [0.00; 0.58] |
| **Total (95% CI)** | | **67** | **10.3%** | **0.06 [0.02; 0.16]** |
| Heterogeneity: $Tau^2$ = 0; $Chi^2$ = 0.88, df = 1 (P = 0.35); $I^2$ = 0% | | | | |
| **Hôtes = Ovis aries** | | | | |
| Chitimia-Dobler et al., 2019 | 1 | 1 | 3.7% | 1.00 [0.02; 1.00] |
| Encinas Grandes, 1986 | 22 | 141 | 6.0% | 0.16 [0.10; 0.23] |
| Dumitrache et al., 2012 | 1 | 5 | 4.6% | 0.20 [0.01; 0.72] |
| **Total (95% CI)** | | **147** | **14.3%** | **0.16 [0.11; 0.23]** |
| Heterogeneity: $Tau^2$ = < 0.0001; $Chi^2$ = 2.9, df = 2 (P = 0.23); $I^2$ = 31% | | | | |
| **Hôtes = Sus scrofa** | | | | |
| Encinas Grandes, 1986 | 1 | 12 | 4.8% | 0.08 [0.00; 0.38] |
| Grassi et al., 2021 | 1 | 55 | 4.9% | 0.02 [0.00; 0.10] |
| **Total (95% CI)** | | **67** | **9.6%** | **0.04 [0.01; 0.14]** |
| Heterogeneity: $Tau^2$ = < 0.0001; $Chi^2$ = 1.2, df = 1 (P = 0.27); $I^2$ = 17% | | | | |
| **Total (95% CI)** | | **3434** | **100.0%** | **0.20 [0.09; 0.40]** |
| Heterogeneity: $Tau^2$ = 3.9838; $Chi^2$ = 171.31, df = 20 (P < 0.01); $I^2$ = 88% | | | | |
| Test for subgroup differences: $Chi^2$ = 98.44, df = 6 (P < 0.01) | | | | |

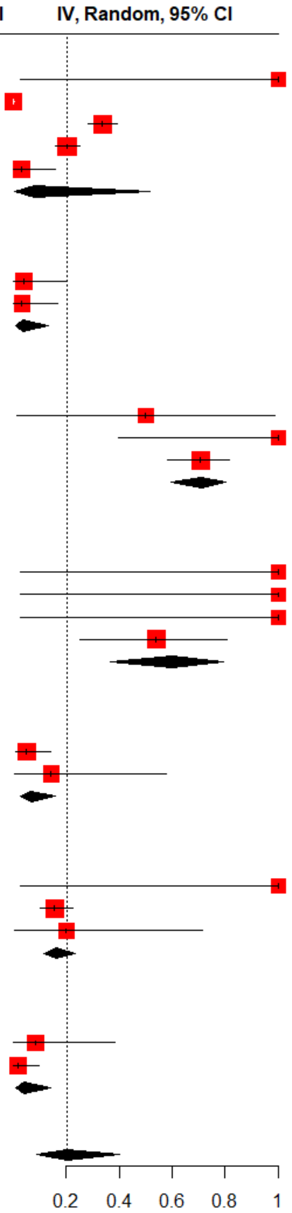


***Figure S1***. *subgroup meta analysis of adult stage infestation rate using host family/species as the grouping variable*

| Study or Subgroup | Events | Total | Weight | IV, Random, 95% CI |
|---|---|---|---|---|
| **Hôtes = Bos taurus** | | | | |
| Cicculli et al., 2019 | 91 | 586 | 2.5% | 0.16 [0.13; 0.19] |
| Torina et al., 2006 | 205 | 3855 | 2.5% | 0.05 [0.05; 0.06] |
| Duzlu et al., 2016 | 245 | 1115 | 2.5% | 0.22 [0.20; 0.25] |
| Papadopoulos et al., 1996 | 409 | 5137 | 2.5% | 0.08 [0.07; 0.09] |
| Yesilbag et al., 2013 | 114 | 501 | 2.5% | 0.23 [0.19; 0.27] |
| Ros-Garcia et al., 2012 | 14 | 94 | 2.5% | 0.15 [0.08; 0.24] |
| Deger et al., 2016 | 36 | 1832 | 2.5% | 0.02 [0.01; 0.03] |
| Aktas et al., 2004 | 3 | 4581 | 2.4% | 0.00 [0.00; 0.00] |
| Encinas Grandes, 1986 | 203 | 1878 | 2.5% | 0.11 [0.09; 0.12] |
| Ica et al., 2007 | 245 | 1160 | 2.5% | 0.21 [0.19; 0.24] |
| Toma et al., 2017 | 49 | 422 | 2.5% | 0.12 [0.09; 0.15] |
| Grech-Angelini et al., 2020 | 417 | 1938 | 2.5% | 0.22 [0.20; 0.23] |
| Bakirci et al., 2011 | 7358 | 19679 | 2.5% | 0.37 [0.37; 0.38] |
| Pereira et al., 2018 | 107 | 147 | 2.5% | 0.73 [0.65; 0.80] |
| Gunes et al., 2011 | 855 | 1294 | 2.5% | 0.66 [0.63; 0.69] |
| Incl et al., 2008 | 272 | 1585 | 2.5% | 0.17 [0.15; 0.19] |
| Le Riche et al., 1974 | 96 | 1802 | 2.5% | 0.05 [0.04; 0.06] |
| Rinaldi et al., 2014 | 170 | 585 | 2.5% | 0.29 [0.25; 0.33] |
| Ica et al., 2007 | 272 | 1585 | 2.5% | 0.17 [0.15; 0.19] |
| **Total (95% CI)** | | 49776 | 47.8% | 0.14 [0.07; 0.25] |
| Heterogeneity: $Tau^2 = 2.5830$; $Chi^2 = 4658.71$, df = 18 (P = 0); $I^2 = 100\%$ | | | | |
| **Hôtes = Equus ferus caballus** | | | | |
| Butler et al., 2016 | 2 | 130 | 2.3% | 0.02 [0.00; 0.05] |
| Duscher et al., 2018 | 1 | 1 | 1.8% | 1.00 [0.02; 1.00] |
| Chisu et al., 2014 | 12 | 12 | 1.9% | 1.00 [0.74; 1.00] |
| Grech-Angelini et al., 2020 | 140 | 180 | 2.5% | 0.78 [0.71; 0.84] |
| **Total (95% CI)** | | 323 | 8.5% | 0.55 [0.07; 0.95] |
| Heterogeneity: $Tau^2 = 6.8349$; $Chi^2 = 56.73$, df = 3 (P < 0.01); $I^2 = 95\%$ | | | | |
| **Hôtes = Ovis aries** | | | | |
| Chaligiannis et al., 2016 | 3 | 1339 | 2.4% | 0.00 [0.00; 0.01] |
| Torina et al., 2006 | 2 | 126 | 2.3% | 0.02 [0.00; 0.06] |
| Pereira et al., 2018 | 26 | 206 | 2.5% | 0.13 [0.08; 0.18] |
| Grech-Angelini et al., 2020 | 4 | 58 | 2.4% | 0.07 [0.02; 0.17] |
| Encinas Grandes, 1986 | 104 | 1012 | 2.5% | 0.10 [0.08; 0.12] |
| Rinaldi et al., 2014 | 19 | 1594 | 2.5% | 0.01 [0.01; 0.02] |
| Gunes et al., 2011 | 203 | 674 | 2.5% | 0.30 [0.27; 0.34] |
| Le Riche et al., 1974 | 171 | 7026 | 2.5% | 0.02 [0.02; 0.03] |
| Papadopoulos et al., 1996 | 1 | 2265 | 2.2% | 0.00 [0.00; 0.00] |
| **Total (95% CI)** | | 14300 | 21.9% | 0.03 [0.01; 0.09] |
| Heterogeneity: $Tau^2 = 3.4943$; $Chi^2 = 783.4$, df = 8 (P < 0.01); $I^2 = 99\%$ | | | | |
| **Hôtes = Capra hircus** | | | | |
| Chaligiannis et al., 2016 | 2 | 769 | 2.3% | 0.00 [0.00; 0.01] |
| Le Riche et al., 1974 | 10 | 5290 | 2.5% | 0.00 [0.00; 0.00] |
| Papadopoulos et al., 1996 | 2 | 3231 | 2.3% | 0.00 [0.00; 0.00] |
| **Total (95% CI)** | | 9290 | 7.2% | 0.00 [0.00; 0.00] |
| Heterogeneity: $Tau^2 = < 0.0001$; $Chi^2 = 2.52$, df = 2 (P = 0.28); $I^2 = 21\%$ | | | | |
| **Hôtes = Sus scrofa** | | | | |
| Pereira et al., 2018 | 1 | 20 | 2.2% | 0.05 [0.00; 0.25] |
| Ruiz-Fons et al., 2006 | 1288 | 1875 | 2.5% | 0.69 [0.67; 0.71] |
| Grech-Angelini et al., 2020 | 8 | 297 | 2.5% | 0.03 [0.01; 0.05] |
| **Total (95% CI)** | | 2192 | 7.2% | 0.14 [0.02; 0.62] |
| Heterogeneity: $Tau^2 = 3.8095$; $Chi^2 = 158.64$, df = 2 (P < 0.01); $I^2 = 99\%$ | | | | |
| **Hôtes = Cervidae** | | | | |
| Keysary et al., 2011 | 14 | 109 | 2.5% | 0.13 [0.07; 0.21] |
| Ruiz-Fons et al., 2006 | 2842 | 4461 | 2.5% | 0.64 [0.62; 0.65] |
| De La Fuente et al., 2004 | 558 | 582 | 2.5% | 0.96 [0.94; 0.97] |
| **Total (95% CI)** | | 5152 | 7.5% | 0.65 [0.15; 0.95] |
| Heterogeneity: $Tau^2 = 4.1868$; $Chi^2 = 227.69$, df = 2 (P < 0.01); $I^2 = 99\%$ | | | | |
| **Total (95% CI)** | | **81033** | **100.0%** | **0.10 [0.05; 0.19]** |
| Heterogeneity: $Tau^2 = 6.0100$; $Chi^2 = 10651.62$, df = 40 (P = 0); $I^2 = 100\%$ | | | | |
| Test for subgroup differences: $Chi^2 = 138.30$, df = 5 (P < 0.01) | | | | |

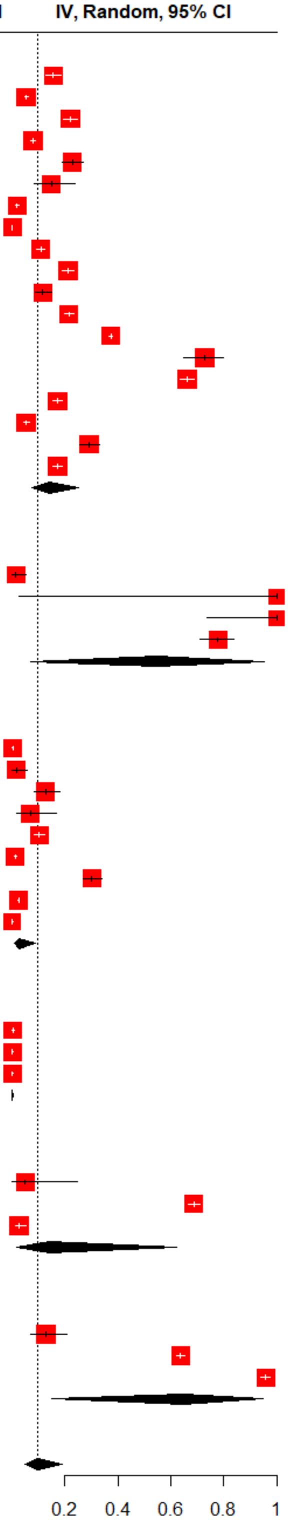


***Figure S2.*** *subgroup meta analysis of adult stage competitivity using host family/species as the grouping variable*

#### Immatures

| Study or Subgroup | Events | Total | Weight | IV, Random, 95% CI |
|---|---|---|---|---|
| **Hôtes = Motacillidae** | | | | |
| Papadopoulos et al., 2001 | 2 | 4 | 1.4% | 0.50 [0.07; 0.93] |
| Kaiser et al., 1974 | 13 | 13 | 1.1% | 1.00 [0.75; 1.00] |
| Kaiser et al., 1974 | 3 | 4 | 1.3% | 0.75 [0.19; 0.99] |
| Movila et al., 2013 | 2 | 2 | 1.1% | 1.00 [0.16; 1.00] |
| Kaiser et al., 1974 | 2 | 2 | 1.1% | 1.00 [0.16; 1.00] |
| **Total (95% CI)** | | **25** | **5.9%** | **0.81 [0.37; 0.97]** |
| Heterogeneity: $Tau^2 = 3.2905$; $Chi^2 = 3.7$, df = 4 (P = 0.45); $I^2 = 0\%$ | | | | |
| **Hôtes = Muscicapidae** | | | | |
| Pereira et al., 2018 | 2 | 2 | 1.1% | 1.00 [0.16; 1.00] |
| Movila et al., 2013 | 2 | 6 | 1.4% | 0.33 [0.04; 0.78] |
| Keskin and Erciyas-Yavuz, 2019 | 17 | 48 | 1.7% | 0.35 [0.22; 0.51] |
| Kaiser et al., 1974 | 9 | 16 | 1.6% | 0.56 [0.30; 0.80] |
| Kaiser et al., 1974 | 23 | 23 | 1.1% | 1.00 [0.85; 1.00] |
| Kaiser et al., 1974 | 1 | 4 | 1.3% | 0.25 [0.01; 0.81] |
| Kaiser et al., 1974 | 11 | 12 | 1.3% | 0.92 [0.62; 1.00] |
| Keskin and Erciyas-Yavuz, 2019 | 1 | 17 | 1.3% | 0.06 [0.00; 0.29] |
| Jameson et al., 2012 | 1 | 2 | 1.1% | 0.50 [0.01; 0.99] |
| Keskin and Erciyas-Yavuz, 2019 | 18 | 21 | 1.5% | 0.86 [0.64; 0.97] |
| Keskin and Erciyas-Yavuz, 2019 | 2 | 2 | 1.1% | 1.00 [0.16; 1.00] |
| Palomar et al., 2015 | 5 | 11 | 1.6% | 0.45 [0.17; 0.77] |
| Kaiser et al., 1974 | 2 | 3 | 1.2% | 0.67 [0.09; 0.99] |
| Keskin and Erciyas-Yavuz, 2019 | 2 | 7 | 1.4% | 0.29 [0.04; 0.71] |
| Jameson et al., 2012 | 8 | 8 | 1.1% | 1.00 [0.63; 1.00] |
| Kaiser et al., 1974 | 5 | 5 | 1.1% | 1.00 [0.48; 1.00] |
| Santos-Silva et al., 2006 | 28 | 28 | 1.1% | 1.00 [0.88; 1.00] |
| Kaiser et al., 1974 | 5 | 5 | 1.1% | 1.00 [0.48; 1.00] |
| Hornok et al., 2013 | 3 | 184 | 1.6% | 0.02 [0.00; 0.05] |
| Keskin and Erciyas-Yavuz, 2019 | 4 | 4 | 1.1% | 1.00 [0.40; 1.00] |
| Keskin and Erciyas-Yavuz, 2019 | 2 | 473 | 1.5% | 0.00 [0.00; 0.02] |
| Papadopoulos et al., 2001 | 12 | 14 | 1.5% | 0.86 [0.57; 0.98] |
| Papadopoulos et al., 2001 | 2 | 26 | 1.5% | 0.08 [0.01; 0.25] |
| Papadopoulos et al., 2001 | 14 | 14 | 1.1% | 1.00 [0.77; 1.00] |
| **Total (95% CI)** | | **935** | **31.6%** | **0.57 [0.36; 0.75]** |
| Heterogeneity: $Tau^2 = 3.2905$; $Chi^2 = 180.32$, df = 23 ($P < 0.01$); $I^2 = 87\%$ | | | | |
| **Hôtes = Laridae** | | | | |
| Pereira et al., 2018 | 5 | 6 | 1.3% | 0.83 [0.36; 1.00] |
| **Hôtes = Strigidae** | | | | |
| Santos-Silva et al., 2006 | 4 | 4 | 1.1% | 1.00 [0.40; 1.00] |
| Pereira et al., 2018 | 1 | 2 | 1.1% | 0.50 [0.01; 0.99] |
| Santos-Silva et al., 2006 | 3 | 3 | 1.1% | 1.00 [0.29; 1.00] |
| Kaiser et al., 1974 | 7 | 7 | 1.1% | 1.00 [0.59; 1.00] |
| Pereira et al., 2018 | 19 | 22 | 1.5% | 0.86 [0.65; 0.97] |
| **Total (95% CI)** | | **38** | **6.0%** | **0.85 [0.45; 0.98]** |
| Heterogeneity: $Tau^2 = 3.2905$; $Chi^2 = 2.09$, df = 4 (P = 0.72); $I^2 = 0\%$ | | | | |
| **Hôtes = Accipitridae** | | | | |
| Santos-Silva et al., 2006 | 7 | 7 | 1.1% | 1.00 [0.59; 1.00] |
| Santos-Silva et al., 2006 | 4 | 4 | 1.1% | 1.00 [0.40; 1.00] |
| **Total (95% CI)** | | **11** | **2.2%** | **0.92 [0.31; 1.00]** |
| Heterogeneity: $Tau^2 = 3.2905$; $Chi^2 = 0.06$, df = 1 (P = 0.81); $I^2 = 0\%$ | | | | |
| **Hôtes = Paridae** | | | | |
| Santos-Silva et al., 2006 | 3 | 3 | 1.1% | 1.00 [0.29; 1.00] |
| Santos-Silva et al., 2006 | 3 | 3 | 1.1% | 1.00 [0.29; 1.00] |
| Palomar et al., 2015 | 3 | 4 | 1.3% | 0.75 [0.19; 0.99] |
| Palomar et al., 2015 | 6 | 8 | 1.5% | 0.75 [0.35; 0.97] |
| **Total (95% CI)** | | **18** | **4.9%** | **0.81 [0.33; 0.97]** |
| Heterogeneity: $Tau^2 = 3.2905$; $Chi^2 = 0.45$, df = 3 (P = 0.93); $I^2 = 0\%$ | | | | |
| **Hôtes = Corvidae** | | | | |
| Norte et al., 2012 | 1 | 37 | 1.4% | 0.03 [0.00; 0.14] |
| Sandor et al., 2017 | 1 | 23 | 1.4% | 0.04 [0.00; 0.22] |
| **Total (95% CI)** | | **60** | **2.7%** | **0.03 [0.00; 0.39]** |
| Heterogeneity: $Tau^2 = 3.2905$; $Chi^2 = 0.12$, df = 1 (P = 0.73); $I^2 = 0\%$ | | | | |

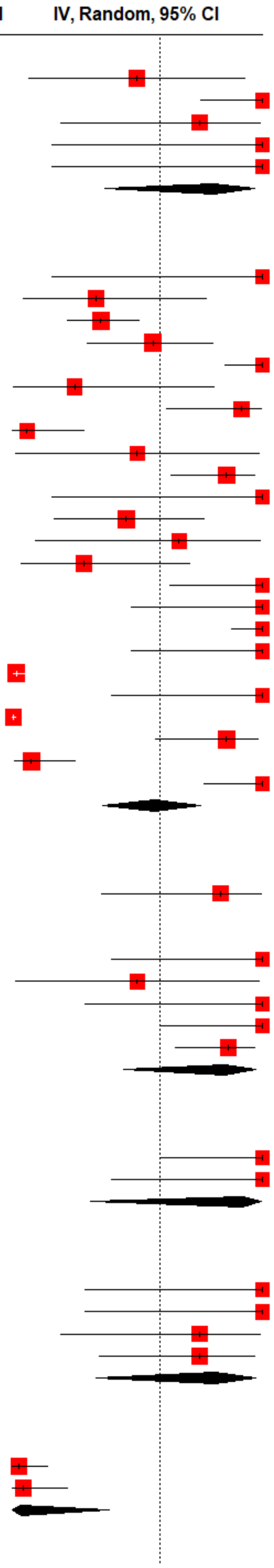

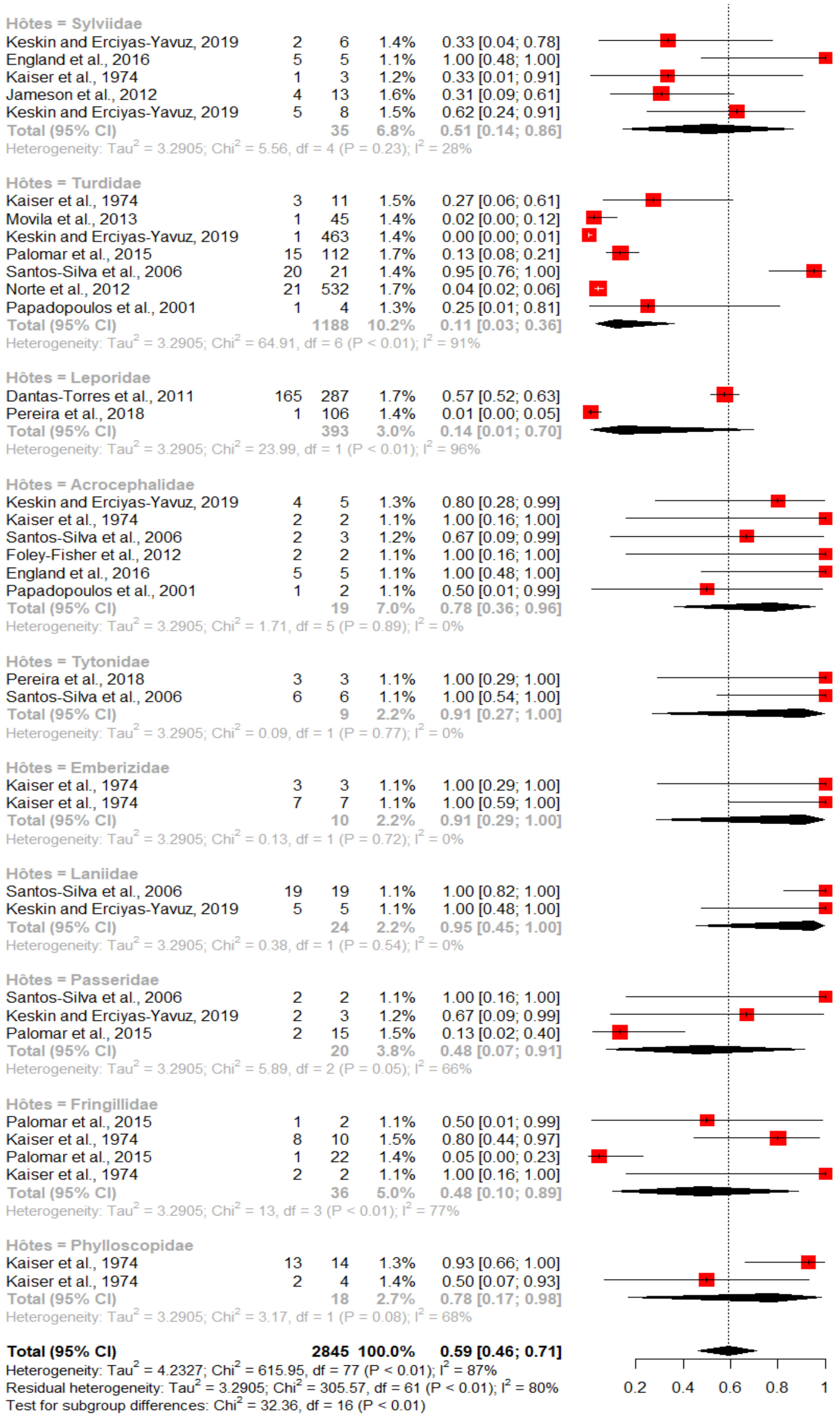


***Figure S3****. subgroup meta analysis of immatures stages competitivity using host family/species as the grouping variable*

| Study or Subgroup | Mean | SD | Total | Weight | Mean IV, Random, 95% CI |
|---|---|---|---|---|---|
| **Hôtes = Phylloscopidae** | | | | | |
| England et al., 2016 | 0.10 | 0.3162 | 10 | 1.3% | 0.10 [-0.10; 0.30] |
| Kaiser et al., 1974 | 0.04 | 0.2099 | 295 | 1.3% | 0.04 [ 0.02; 0.07] |
| Kaiser et al., 1974 | 0.05 | 0.2209 | 41 | 1.3% | 0.05 [-0.02; 0.12] |
| **Total (95% CI)** | | | 346 | 3.9% | 0.06 [-1.03; 1.16] |
| Heterogeneity: $Tau^2 = 0.9280$; $Chi^2 = 0.32$, df = 2 (P = 0.85); $I^2 = 0\%$ | | | | | |
| **Hôtes = Acrocephalidae** | | | | | |
| Jameson et al., 2012 | 0.10 | 0.3162 | 10 | 1.3% | 0.10 [-0.10; 0.30] |
| Kaiser et al., 1974 | 0.07 | 0.2673 | 14 | 1.3% | 0.07 [-0.07; 0.21] |
| Keskin and Erciyas-Yavuz, 2019 | 0.21 | 0.4588 | 19 | 1.3% | 0.21 [ 0.00; 0.42] |
| Kaiser et al., 1974 | 0.08 | 0.2887 | 24 | 1.3% | 0.08 [-0.03; 0.20] |
| Santos-Silva et al., 2006 | 0.67 | 0.8165 | 3 | 1.1% | 0.67 [-0.26; 1.59] |
| Capek et al., 2014 | 0.01 | 0.1109 | 325 | 1.3% | 0.01 [ 0.00; 0.02] |
| Capek et al., 2014 | 0.03 | 0.1733 | 333 | 1.3% | 0.03 [ 0.01; 0.05] |
| Palomar et al., 2016 | 2.00 | 1.4142 | 5 | 1.0% | 2.00 [ 0.76; 3.24] |
| Foley-Fisher et al., 2012 | 0.25 | 0.5000 | 8 | 1.3% | 0.25 [-0.10; 0.60] |
| England et al., 2016 | 0.62 | 0.7906 | 8 | 1.2% | 0.62 [ 0.08; 1.17] |
| England et al., 2016 | 0.12 | 0.3536 | 8 | 1.3% | 0.12 [-0.12; 0.37] |
| Capek et al., 2014 | 0.11 | 0.3273 | 84 | 1.3% | 0.11 [ 0.04; 0.18] |
| **Total (95% CI)** | | | 841 | 15.1% | 0.31 [-0.25; 0.87] |
| Heterogeneity: $Tau^2 = 0.9280$; $Chi^2 = 32.9$, df = 11 (P < 0.01); $I^2 = 67\%$ | | | | | |
| **Hôtes = Sylviidae** | | | | | |
| Keskin and Erciyas-Yavuz, 2019 | 0.00 | 0.0446 | 1004 | 1.3% | 0.00 [-0.00; 0.00] |
| England et al., 2016 | 0.42 | 0.6455 | 12 | 1.3% | 0.42 [ 0.05; 0.78] |
| Kaiser et al., 1974 | 0.05 | 0.2236 | 20 | 1.3% | 0.05 [-0.05; 0.15] |
| Kaiser et al., 1974 | 0.00 | 0.0626 | 255 | 1.3% | 0.00 [-0.00; 0.01] |
| Jameson et al., 2012 | 0.08 | 0.2747 | 53 | 1.3% | 0.08 [ 0.00; 0.15] |
| Keskin and Erciyas-Yavuz, 2019 | 0.01 | 0.0762 | 860 | 1.3% | 0.01 [ 0.00; 0.01] |
| **Total (95% CI)** | | | 2204 | 7.9% | 0.09 [-0.68; 0.86] |
| Heterogeneity: $Tau^2 = 0.9280$; $Chi^2 = 11.24$, df = 5 (P = 0.05); $I^2 = 55\%$ | | | | | |
| **Hôtes = Muscicapidae** | | | | | |
| Movila et al., 2013 | 0.18 | 0.4264 | 11 | 1.3% | 0.18 [-0.07; 0.43] |
| Keskin and Erciyas-Yavuz, 2019 | 0.15 | 0.3862 | 114 | 1.3% | 0.15 [ 0.08; 0.22] |
| Kaiser et al., 1974 | 0.07 | 0.2554 | 138 | 1.3% | 0.07 [ 0.02; 0.11] |
| Kaiser et al., 1974 | 1.64 | 1.2817 | 14 | 1.2% | 1.64 [ 0.97; 2.31] |
| Kaiser et al., 1974 | 0.06 | 0.2425 | 17 | 1.3% | 0.06 [-0.06; 0.17] |
| Kaiser et al., 1974 | 0.41 | 0.6383 | 27 | 1.3% | 0.41 [ 0.17; 0.65] |
| Keskin and Erciyas-Yavuz, 2019 | 0.00 | 0.0579 | 298 | 1.3% | 0.00 [-0.00; 0.01] |
| Jameson et al., 2012 | 0.03 | 0.1826 | 30 | 1.3% | 0.03 [-0.03; 0.10] |
| Keskin and Erciyas-Yavuz, 2019 | 0.58 | 0.7620 | 31 | 1.3% | 0.58 [ 0.31; 0.85] |
| Keskin and Erciyas-Yavuz, 2019 | 0.06 | 0.2425 | 34 | 1.3% | 0.06 [-0.02; 0.14] |
| Palomar et al., 2016 | 2.50 | 1.5811 | 4 | 0.8% | 2.50 [ 0.95; 4.05] |
| Palomar et al., 2015 | 1.25 | 1.1180 | 4 | 1.0% | 1.25 [ 0.15; 2.35] |
| Kaiser et al., 1974 | 0.05 | 0.2182 | 42 | 1.3% | 0.05 [-0.02; 0.11] |
| Keskin and Erciyas-Yavuz, 2019 | 0.00 | 0.0640 | 489 | 1.3% | 0.00 [-0.00; 0.01] |
| Jameson et al., 2012 | 0.16 | 0.3961 | 51 | 1.3% | 0.16 [ 0.05; 0.27] |
| Kaiser et al., 1974 | 0.09 | 0.2936 | 58 | 1.3% | 0.09 [ 0.01; 0.16] |
| Santos-Silva et al., 2006 | 4.67 | 2.1602 | 6 | 0.8% | 4.67 [ 2.94; 6.40] |
| Kaiser et al., 1974 | 0.71 | 0.8452 | 7 | 1.2% | 0.71 [ 0.09; 1.34] |
| Kaiser et al., 1974 | 0.14 | 0.3780 | 7 | 1.3% | 0.14 [-0.14; 0.42] |
| Hornok et al., 2013 | 0.00 | 0.0639 | 734 | 1.3% | 0.00 [-0.00; 0.01] |
| Keskin and Erciyas-Yavuz, 2019 | 0.44 | 0.6667 | 9 | 1.3% | 0.44 [ 0.01; 0.88] |
| Keskin and Erciyas-Yavuz, 2019 | 0.00 | 0.0471 | 903 | 1.3% | 0.00 [-0.00; 0.01] |
| **Total (95% CI)** | | | 3028 | 27.3% | 0.46 [ 0.04; 0.88] |
| Heterogeneity: $Tau^2 = 0.9280$; $Chi^2 = 148.57$, df = 21 (P < 0.01); $I^2 = 86\%$ | | | | | |

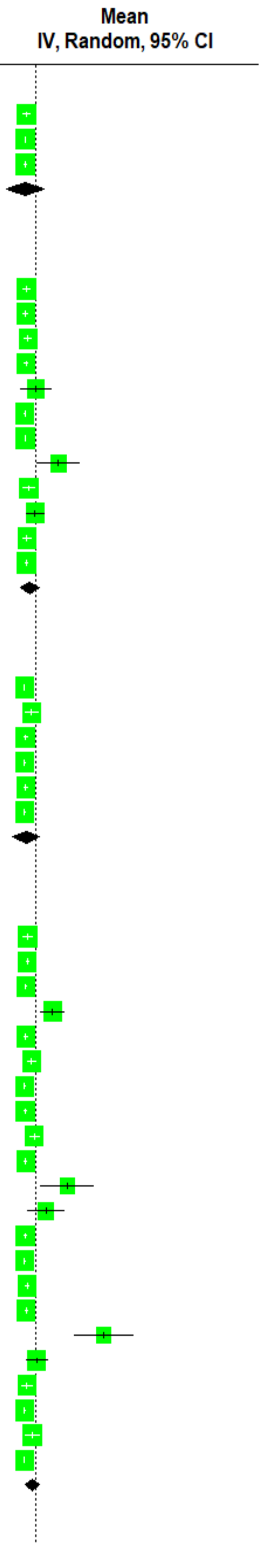

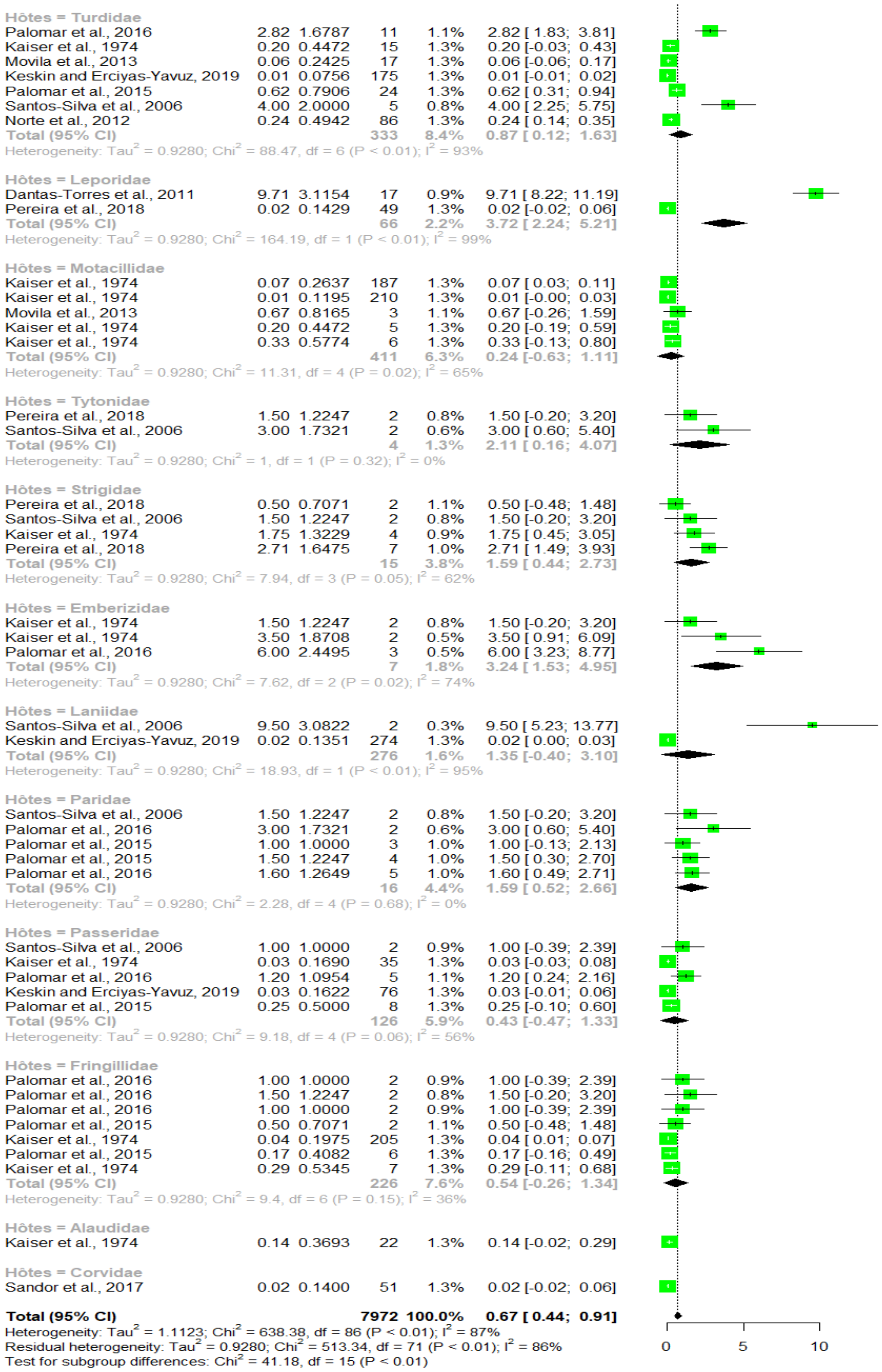


***Figure S4.*** *subgroup meta analysis of immatures stages parasitic load using host family/species as the grouping variable*

### Landscape

#### Immatures

| Study or Subgroup | Events | Total | Weight (common) | Weight (random) | IV, Fixed + Random, 95% CI |
|---|---|---|---|---|---|
| Landscape = Mediterranean | | | | | |
| Pereira et al., 2018 | 1 | 1 | 0.1% | 1.2% | 1.00 [0.02; 1.00] |
| Pereira et al., 2018 | 1 | 1 | 0.1% | 1.2% | 1.00 [0.02; 1.00] |
| Pereira et al., 2018 | 1 | 1 | 0.1% | 1.2% | 1.00 [0.02; 1.00] |
| Pereira et al., 2018 | 1 | 1 | 0.1% | 1.2% | 1.00 [0.02; 1.00] |
| Pereira et al., 2018 | 1 | 1 | 0.1% | 1.2% | 1.00 [0.02; 1.00] |
| Pereira et al., 2018 | 1 | 1 | 0.1% | 1.2% | 1.00 [0.02; 1.00] |
| Encinas Grandes, 1986 | 1 | 1 | 0.1% | 1.2% | 1.00 [0.02; 1.00] |
| Encinas Grandes, 1986 | 1 | 1 | 0.1% | 1.2% | 1.00 [0.02; 1.00] |
| Encinas Grandes, 1986 | 1 | 1 | 0.1% | 1.2% | 1.00 [0.02; 1.00] |
| Santos-Silva et al., 2006 | 1 | 1 | 0.1% | 1.2% | 1.00 [0.02; 1.00] |
| Santos-Silva et al., 2006 | 1 | 1 | 0.1% | 1.2% | 1.00 [0.02; 1.00] |
| Santos-Silva et al., 2006 | 1 | 1 | 0.1% | 1.2% | 1.00 [0.02; 1.00] |
| Santos-Silva et al., 2006 | 1 | 1 | 0.1% | 1.2% | 1.00 [0.02; 1.00] |
| Santos-Silva et al., 2006 | 1 | 1 | 0.1% | 1.2% | 1.00 [0.02; 1.00] |
| Santos-Silva et al., 2006 | 1 | 1 | 0.1% | 1.2% | 1.00 [0.02; 1.00] |
| Norte et al., 2012 | 1 | 1 | 0.1% | 1.2% | 1.00 [0.02; 1.00] |
| Palomar et al., 2016 | 1 | 1 | 0.1% | 1.2% | 1.00 [0.02; 1.00] |
| Palomar et al., 2016 | 1 | 1 | 0.1% | 1.2% | 1.00 [0.02; 1.00] |
| Palomar et al., 2016 | 1 | 1 | 0.1% | 1.2% | 1.00 [0.02; 1.00] |
| Palomar et al., 2016 | 1 | 1 | 0.1% | 1.2% | 1.00 [0.02; 1.00] |
| Palomar et al., 2016 | 1 | 1 | 0.1% | 1.2% | 1.00 [0.02; 1.00] |
| Palomar et al., 2016 | 1 | 1 | 0.1% | 1.2% | 1.00 [0.02; 1.00] |
| Palomar et al., 2016 | 1 | 1 | 0.1% | 1.2% | 1.00 [0.02; 1.00] |
| Palomar et al., 2016 | 1 | 1 | 0.1% | 1.2% | 1.00 [0.02; 1.00] |
| Palomar et al., 2016 | 1 | 1 | 0.1% | 1.2% | 1.00 [0.02; 1.00] |
| Palomar et al., 2016 | 1 | 1 | 0.1% | 1.2% | 1.00 [0.02; 1.00] |
| Palomar et al., 2015 | 1 | 1 | 0.1% | 1.2% | 1.00 [0.02; 1.00] |
| England et al., 2016 | 1 | 10 | 0.3% | 1.5% | 0.10 [0.00; 0.45] |
| Palomar et al., 2016 | 11 | 11 | 0.2% | 1.3% | 1.00 [0.72; 1.00] |
| Encinas Grandes, 1986 | 8 | 12 | 0.9% | 1.7% | 0.67 [0.35; 0.90] |
| England et al., 2016 | 2 | 12 | 0.6% | 1.7% | 0.17 [0.02; 0.48] |
| Santos-Silva et al., 2006 | 2 | 2 | 0.1% | 1.3% | 1.00 [0.16; 1.00] |
| Santos-Silva et al., 2006 | 2 | 2 | 0.1% | 1.3% | 1.00 [0.16; 1.00] |
| Santos-Silva et al., 2006 | 2 | 2 | 0.1% | 1.3% | 1.00 [0.16; 1.00] |
| Santos-Silva et al., 2006 | 2 | 2 | 0.1% | 1.3% | 1.00 [0.16; 1.00] |
| Santos-Silva et al., 2006 | 2 | 2 | 0.1% | 1.3% | 1.00 [0.16; 1.00] |
| Santos-Silva et al., 2006 | 2 | 2 | 0.1% | 1.3% | 1.00 [0.16; 1.00] |
| Palomar et al., 2016 | 2 | 2 | 0.1% | 1.3% | 1.00 [0.16; 1.00] |
| Palomar et al., 2016 | 2 | 2 | 0.1% | 1.3% | 1.00 [0.16; 1.00] |
| Palomar et al., 2016 | 2 | 2 | 0.1% | 1.3% | 1.00 [0.16; 1.00] |
| Palomar et al., 2016 | 2 | 2 | 0.1% | 1.3% | 1.00 [0.16; 1.00] |
| Encinas Grandes, 1986 | 3 | 3 | 0.2% | 1.3% | 1.00 [0.29; 1.00] |
| Encinas Grandes, 1986 | 1 | 3 | 0.2% | 1.4% | 0.33 [0.01; 0.91] |
| Santos-Silva et al., 2006 | 3 | 3 | 0.2% | 1.3% | 1.00 [0.29; 1.00] |
| Palomar et al., 2016 | 3 | 3 | 0.2% | 1.3% | 1.00 [0.29; 1.00] |
| Palomar et al., 2016 | 4 | 4 | 0.2% | 1.3% | 1.00 [0.40; 1.00] |
| Pereira et al., 2018 | 1 | 49 | 0.3% | 1.5% | 0.02 [0.00; 0.11] |
| Santos-Silva et al., 2006 | 5 | 5 | 0.2% | 1.3% | 1.00 [0.48; 1.00] |
| Palomar et al., 2016 | 5 | 5 | 0.2% | 1.3% | 1.00 [0.48; 1.00] |
| Palomar et al., 2016 | 5 | 5 | 0.2% | 1.3% | 1.00 [0.48; 1.00] |
| Palomar et al., 2016 | 5 | 5 | 0.2% | 1.3% | 1.00 [0.48; 1.00] |
| Santos-Silva et al., 2006 | 6 | 6 | 0.2% | 1.3% | 1.00 [0.54; 1.00] |
| Hoffman et al., 2020 | 203 | 7354 | 68.5% | 1.9% | 0.03 [0.02; 0.03] |
| Foley-Fisher et al., 2012 | 1 | 8 | 0.3% | 1.5% | 0.12 [0.00; 0.53] |
| England et al., 2016 | 1 | 8 | 0.3% | 1.5% | 0.12 [0.00; 0.53] |
| England et al., 2016 | 1 | 8 | 0.3% | 1.5% | 0.12 [0.00; 0.53] |
| Total (common effect, 95% CI) | | 7561 | 78.3% | -- | 0.05 [0.04; 0.05] |
| Total (random effect, 95% CI) | | | -- | 72.3% | 0.66 [0.54; 0.77] |
| Heterogeneity: $Tau^2 = 1.8700$; $Chi^2 = 497.99$, df = 55 ($P < 0.01$); $I^2 = 89\%$ | | | | | |
| Landscape = Continental | | | | | |
| Kolodziejek et al., 2014 | 1 | 1 | 0.1% | 1.2% | 1.00 [0.02; 1.00] |
| Capek et al., 2014 | 2 | 325 | 0.7% | 1.7% | 0.01 [0.00; 0.02] |
| Capek et al., 2014 | 7 | 333 | 2.4% | 1.8% | 0.02 [0.01; 0.04] |
| Sandor et al., 2017 | 1 | 51 | 0.3% | 1.5% | 0.02 [0.00; 0.10] |
| Hornok et al., 2013 | 42 | 734 | 13.7% | 1.9% | 0.06 [0.04; 0.08] |
| Capek et al., 2014 | 1 | 84 | 0.3% | 1.5% | 0.01 [0.00; 0.06] |
| Capek et al., 2014 | 1 | 90 | 0.3% | 1.5% | 0.01 [0.00; 0.06] |
| Total (common effect, 95% CI) | | 1618 | 18.0% | -- | 0.04 [0.03; 0.06] |
| Total (random effect, 95% CI) | | | -- | 11.2% | 0.03 [0.01; 0.08] |
| Heterogeneity: $Tau^2 = 1.8700$; $Chi^2 = 25.98$, df = 6 ($P < 0.01$); $I^2 = 77\%$ | | | | | |
| Landscape = Steppic | | | | | |
| Palomar et al., 2016 | 1 | 1 | 0.1% | 1.2% | 1.00 [0.02; 1.00] |
| Dantas-Torres et al., 2011 | 17 | 17 | 0.2% | 1.3% | 1.00 [0.80; 1.00] |
| Total (common effect, 95% CI) | | 18 | 0.3% | -- | 0.92 [0.59; 0.99] |
| Total (random effect, 95% CI) | | | -- | 2.5% | 0.92 [0.39; 0.99] |
| Heterogeneity: $Tau^2 = 1.8700$; $Chi^2 = 1.28$, df = 1 ($P = 0.26$); $I^2 = 22\%$ | | | | | |
| Landscape = Boreal | | | | | |
| Movila et al., 2013 | 1 | 11 | 0.3% | 1.5% | 0.09 [0.00; 0.41] |
| Movila et al., 2013 | 1 | 17 | 0.3% | 1.5% | 0.06 [0.00; 0.29] |
| Movila et al., 2013 | 2 | 3 | 0.2% | 1.4% | 0.67 [0.09; 0.99] |
| Total (common effect, 95% CI) | | 31 | 0.9% | -- | 0.16 [0.05; 0.39] |
| Total (random effect, 95% CI) | | | -- | 4.5% | 0.17 [0.03; 0.61] |
| Heterogeneity: $Tau^2 = 1.8700$; $Chi^2 = 5.23$, df = 2 ($P = 0.07$); $I^2 = 62\%$ | | | | | |
| Landscape = Anatolian | | | | | |
| Keskin and Erciyas-Yavuz, 2019 | 1 | 175 | 0.3% | 1.5% | 0.01 [0.00; 0.03] |
| Keskin and Erciyas-Yavuz, 2019 | 2 | 274 | 0.7% | 1.7% | 0.01 [0.00; 0.03] |
| Keskin and Erciyas-Yavuz, 2019 | 1 | 298 | 0.3% | 1.5% | 0.00 [0.00; 0.02] |
| Keskin and Erciyas-Yavuz, 2019 | 1 | 34 | 0.3% | 1.5% | 0.03 [0.00; 0.15] |
| Keskin and Erciyas-Yavuz, 2019 | 1 | 69 | 0.3% | 1.5% | 0.01 [0.00; 0.08] |
| Keskin and Erciyas-Yavuz, 2019 | 2 | 9 | 0.5% | 1.6% | 0.22 [0.03; 0.60] |
| Total (common effect, 95% CI) | | 859 | 2.6% | -- | 0.02 [0.01; 0.04] |
| Total (random effect, 95% CI) | | | -- | 9.5% | 0.02 [0.00; 0.06] |
| Heterogeneity: $Tau^2 = 1.8700$; $Chi^2 = 17.88$, df = 5 ($P < 0.01$); $I^2 = 72\%$ | | | | | |
| **Total (common effect, 95% CI)** | | **10087** | **100.0%** | **--** | **0.05 [0.04; 0.05]** |
| **Total (random effect, 95% CI)** | | | **--** | **100.0%** | **0.45 [0.31; 0.60]** |

Heterogeneity: $Tau^2 = 4.9484$; $Chi^2 = 586.46$, df = 73 ($P < 0.01$); $I^2 = 88\%$
Residual heterogeneity: $Tau^2 = 1.8700$; $Chi^2 = 548.35$, df = 69 ($P < 0.01$); $I^2 = 87\%$
Test for subgroup differences (common effect): $Chi^2 = 38.10$, df = 4 ($P < 0.01$)
Test for subgroup differences (random effects): $Chi^2 = 79.00$, df = 4 ($P < 0.01$)

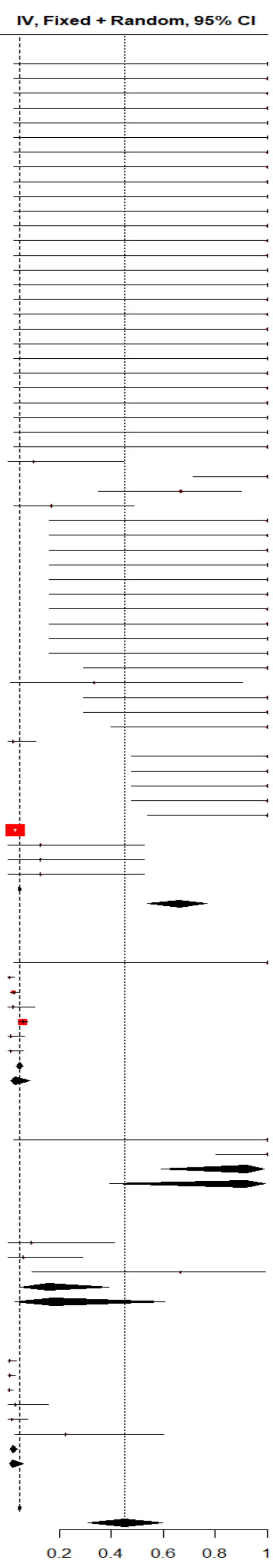


***Figure S5.*** *subgroup meta analysis of immatures stages infestation rate using landscape as the grouping variable*

| Study or Subgroup | Events | Total | Weight (common) | Weight (random) | IV, Fixed + Random, 95% CI |
|---|---|---|---|---|---|
| **Landscape = Alpine** | | | | | |
| Papadopoulos et al., 2001 | 2 | 4 | 0.4% | 1.5% | 0.50 [0.07; 0.93] |
| Hagman et al., 2014 | 659 | 747 | 33.4% | 1.9% | 0.88 [0.86; 0.90] |
| Papadopoulos et al., 2001 | 12 | 14 | 0.7% | 1.7% | 0.86 [0.57; 0.98] |
| Papadopoulos et al., 2001 | 2 | 26 | 0.8% | 1.7% | 0.08 [0.01; 0.25] |
| Papadopoulos et al., 2001 | 14 | 14 | 0.2% | 1.3% | 1.00 [0.77; 1.00] |
| Papadopoulos et al., 2001 | 1 | 4 | 0.3% | 1.4% | 0.25 [0.01; 0.81] |
| Papadopoulos et al., 2001 | 1 | 2 | 0.2% | 1.3% | 0.50 [0.01; 0.99] |
| Total (common effect, 95% CI) | | 811 | 36.1% | -- | 0.86 [0.84; 0.89] |
| Total (random effect, 95% CI) | | | -- | 10.7% | 0.62 [0.23; 0.90] |
| Heterogeneity: $Tau^2$ = 4.1905; $Chi^2$ = 49.54, df = 6 (P < 0.01); $I^2$ = 88% | | | | | |
| **Landscape = Mediterranean** | | | | | |
| Pereira et al., 2018 | 1 | 1 | 0.2% | 1.2% | 1.00 [0.02; 1.00] |
| Pereira et al., 2018 | 2 | 2 | 0.2% | 1.2% | 1.00 [0.16; 1.00] |
| Pereira et al., 2018 | 1 | 1 | 0.2% | 1.2% | 1.00 [0.02; 1.00] |
| Pereira et al., 2018 | 1 | 1 | 0.2% | 1.2% | 1.00 [0.02; 1.00] |
| Pereira et al., 2018 | 1 | 1 | 0.2% | 1.2% | 1.00 [0.02; 1.00] |
| Pereira et al., 2018 | 5 | 6 | 0.4% | 1.5% | 0.83 [0.36; 1.00] |
| Santos-Silva et al., 2006 | 4 | 4 | 0.2% | 1.3% | 1.00 [0.40; 1.00] |
| Santos-Silva et al., 2006 | 1 | 1 | 0.2% | 1.2% | 1.00 [0.02; 1.00] |
| Santos-Silva et al., 2006 | 1 | 1 | 0.2% | 1.2% | 1.00 [0.02; 1.00] |
| Santos-Silva et al., 2006 | 7 | 7 | 0.2% | 1.3% | 1.00 [0.59; 1.00] |
| Santos-Silva et al., 2006 | 4 | 4 | 0.2% | 1.3% | 1.00 [0.40; 1.00] |
| Santos-Silva et al., 2006 | 3 | 3 | 0.2% | 1.2% | 1.00 [0.29; 1.00] |
| Norte et al., 2012 | 1 | 37 | 0.4% | 1.5% | 0.03 [0.00; 0.14] |
| Palomar et al., 2015 | 1 | 1 | 0.2% | 1.2% | 1.00 [0.02; 1.00] |
| England et al., 2016 | 1 | 1 | 0.2% | 1.2% | 1.00 [0.02; 1.00] |
| England et al., 2016 | 5 | 5 | 0.2% | 1.3% | 1.00 [0.48; 1.00] |
| Palomar et al., 2017 | 34 | 336 | 13.2% | 1.8% | 0.10 [0.07; 0.14] |
| Pereira et al., 2018 | 3 | 3 | 0.2% | 1.2% | 1.00 [0.29; 1.00] |
| Pereira et al., 2018 | 1 | 2 | 0.2% | 1.3% | 0.50 [0.01; 0.99] |
| Pereira et al., 2018 | 7 | 7 | 0.2% | 1.3% | 1.00 [0.59; 1.00] |
| Santos-Silva et al., 2006 | 3 | 3 | 0.2% | 1.2% | 1.00 [0.29; 1.00] |
| Santos-Silva et al., 2006 | 19 | 19 | 0.2% | 1.3% | 1.00 [0.82; 1.00] |
| Santos-Silva et al., 2006 | 15 | 15 | 0.2% | 1.3% | 1.00 [0.78; 1.00] |
| Santos-Silva et al., 2006 | 3 | 3 | 0.2% | 1.2% | 1.00 [0.29; 1.00] |
| Santos-Silva et al., 2006 | 2 | 2 | 0.2% | 1.2% | 1.00 [0.16; 1.00] |
| Santos-Silva et al., 2006 | 6 | 6 | 0.2% | 1.3% | 1.00 [0.54; 1.00] |
| Palomar et al., 2015 | 1 | 2 | 0.2% | 1.3% | 0.50 [0.01; 0.99] |
| Palomar et al., 2015 | 15 | 112 | 5.6% | 1.8% | 0.13 [0.08; 0.21] |
| Santos-Silva et al., 2006 | 2 | 3 | 0.3% | 1.4% | 0.67 [0.09; 0.99] |
| Palomar et al., 2015 | 3 | 4 | 0.3% | 1.4% | 0.75 [0.19; 0.99] |
| Palomar et al., 2015 | 6 | 8 | 0.6% | 1.6% | 0.75 [0.35; 0.97] |
| Palomar et al., 2015 | 5 | 11 | 1.2% | 1.7% | 0.45 [0.17; 0.77] |
| Pereira et al., 2018 | 1 | 106 | 0.4% | 1.5% | 0.01 [0.00; 0.05] |
| Santos-Silva et al., 2006 | 20 | 21 | 0.4% | 1.5% | 0.95 [0.76; 1.00] |
| Santos-Silva et al., 2006 | 28 | 28 | 0.2% | 1.3% | 1.00 [0.88; 1.00] |
| Palomar et al., 2015 | 1 | 22 | 0.4% | 1.5% | 0.05 [0.00; 0.23] |
| Pereira et al., 2018 | 19 | 22 | 1.1% | 1.7% | 0.86 [0.65; 0.97] |
| Hoffman et al., 2020 | 331 | 358 | 10.8% | 1.8% | 0.92 [0.89; 0.95] |
| Foley-Fisher et al., 2012 | 2 | 2 | 0.2% | 1.2% | 1.00 [0.16; 1.00] |
| England et al., 2016 | 5 | 5 | 0.2% | 1.3% | 1.00 [0.48; 1.00] |
| England et al., 2016 | 1 | 1 | 0.2% | 1.2% | 1.00 [0.02; 1.00] |
| Palomar et al., 2015 | 2 | 15 | 0.7% | 1.7% | 0.13 [0.02; 0.40] |
| Norte et al., 2012 | 21 | 532 | 8.7% | 1.8% | 0.04 [0.02; 0.06] |
| Total (common effect, 95% CI) | | 1724 | 49.7% | -- | 0.35 [0.31; 0.39] |
| Total (random effect, 95% CI) | | | -- | 59.2% | 0.69 [0.52; 0.82] |
| Heterogeneity: $Tau^2$ = 4.1905; $Chi^2$ = 632.25, df = 42 (P < 0.01); $I^2$ = 93% | | | | | |
| **Landscape = Continental** | | | | | |
| Kolodziejek et al., 2014 | 1 | 1 | 0.2% | 1.2% | 1.00 [0.02; 1.00] |
| Sandor et al., 2017 | 1 | 23 | 0.4% | 1.5% | 0.04 [0.00; 0.22] |
| Hornok et al., 2013 | 3 | 184 | 1.3% | 1.7% | 0.02 [0.00; 0.05] |
| Total (common effect, 95% CI) | | 208 | 1.8% | -- | 0.03 [0.01; 0.08] |
| Total (random effect, 95% CI) | | | -- | 4.4% | 0.08 [0.01; 0.56] |
| Heterogeneity: $Tau^2$ = 4.1905; $Chi^2$ = 9.12, df = 2 (P = 0.01); $I^2$ = 78% | | | | | |
| **Landscape = Anatolian** | | | | | |
| Keskin and Erciyas-Yavuz, 2019 | 2 | 6 | 0.6% | 1.6% | 0.33 [0.04; 0.78] |
| Keskin and Erciyas-Yavuz, 2019 | 17 | 48 | 4.7% | 1.8% | 0.35 [0.22; 0.51] |
| Keskin and Erciyas-Yavuz, 2019 | 1 | 463 | 0.4% | 1.5% | 0.00 [0.00; 0.01] |
| Keskin and Erciyas-Yavuz, 2019 | 4 | 5 | 0.3% | 1.5% | 0.80 [0.28; 0.99] |
| Keskin and Erciyas-Yavuz, 2019 | 5 | 5 | 0.2% | 1.3% | 1.00 [0.48; 1.00] |
| Keskin and Erciyas-Yavuz, 2019 | 1 | 17 | 0.4% | 1.5% | 0.06 [0.00; 0.29] |
| Keskin and Erciyas-Yavuz, 2019 | 18 | 21 | 1.1% | 1.7% | 0.86 [0.64; 0.97] |
| Keskin and Erciyas-Yavuz, 2019 | 2 | 2 | 0.2% | 1.2% | 1.00 [0.16; 1.00] |
| Keskin and Erciyas-Yavuz, 2019 | 2 | 7 | 0.6% | 1.6% | 0.29 [0.04; 0.71] |
| Keskin and Erciyas-Yavuz, 2019 | 1 | 16 | 0.4% | 1.5% | 0.06 [0.00; 0.30] |
| Keskin and Erciyas-Yavuz, 2019 | 2 | 3 | 0.3% | 1.4% | 0.67 [0.09; 0.99] |
| Keskin and Erciyas-Yavuz, 2019 | 5 | 8 | 0.8% | 1.7% | 0.62 [0.24; 0.91] |
| Keskin and Erciyas-Yavuz, 2019 | 4 | 4 | 0.2% | 1.3% | 1.00 [0.40; 1.00] |
| Keskin and Erciyas-Yavuz, 2019 | 2 | 473 | 0.9% | 1.7% | 0.00 [0.00; 0.02] |
| Total (common effect, 95% CI) | | 1078 | 11.1% | -- | 0.31 [0.23; 0.39] |
| Total (random effect, 95% CI) | | | -- | 21.3% | 0.32 [0.12; 0.61] |
| Heterogeneity: $Tau^2$ = 4.1905; $Chi^2$ = 116.18, df = 13 (P < 0.01); $I^2$ = 89% | | | | | |
| **Landscape = Boreal** | | | | | |
| Movila et al., 2013 | 2 | 6 | 0.6% | 1.6% | 0.33 [0.04; 0.78] |
| Movila et al., 2013 | 1 | 45 | 0.4% | 1.5% | 0.02 [0.00; 0.12] |
| Movila et al., 2013 | 2 | 2 | 0.2% | 1.2% | 1.00 [0.16; 1.00] |
| Total (common effect, 95% CI) | | 53 | 1.2% | -- | 0.19 [0.07; 0.43] |
| Total (random effect, 95% CI) | | | -- | 4.4% | 0.24 [0.02; 0.82] |
| Heterogeneity: $Tau^2$ = 4.1905; $Chi^2$ = 9.99, df = 2 (P < 0.01); $I^2$ = 80% | | | | | |
| **Total (common effect, 95% CI)** | | **3874** | **100.0%** | **--** | **0.55 [0.51; 0.58]** |
| **Total (random effect, 95% CI)** | | | **--** | **100.0%** | **0.56 [0.41; 0.69]** |

Heterogeneity: $Tau^2$ = 4.6439; $Chi^2$ = 1217.07, df = 69 (P < 0.01); $I^2$ = 94%
Residual heterogeneity: $Tau^2$ = 4.1905; $Chi^2$ = 817.08, df = 65 (P < 0.01); $I^2$ = 92%
Test for subgroup differences (common effect): $Chi^2$ = 399.99, df = 4 (P < 0.01)
Test for subgroup differences (random effects): $Chi^2$ = 10.01, df = 4 (P = 0.04)

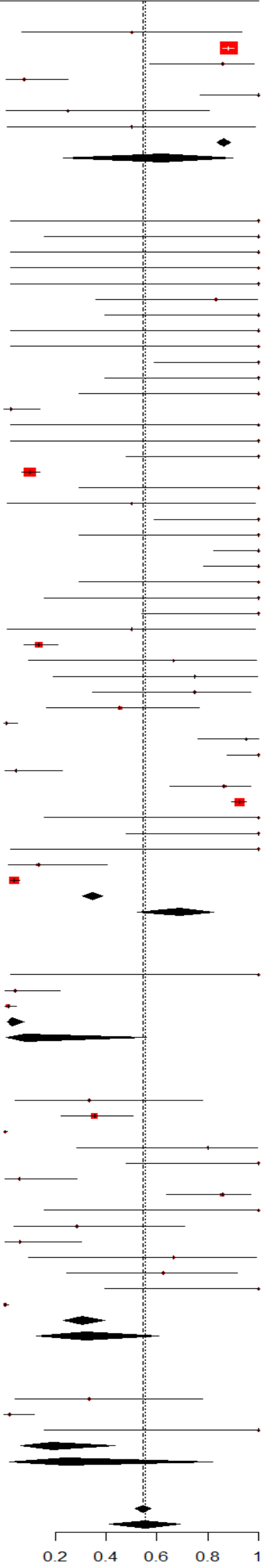


***Figure S6.*** *subgroup meta analysis of immatures stages competitivity using landscape as the grouping variable*

| Study or Subgroup | Mean | SD | Total | Weight | Mean IV, Random, 95% CI |
|---|---|---|---|---|---|
| **Landscape = Mediterranean** | | | | | |
| Pereira et al., 2018 | 1.00 | 1.0000 | 1 | 1.0% | 1.00 [-0.96; 2.96] |
| Pereira et al., 2018 | 2.00 | 1.4142 | 1 | 0.7% | 2.00 [-0.77; 4.77] |
| Pereira et al., 2018 | 1.00 | 1.0000 | 1 | 1.0% | 1.00 [-0.96; 2.96] |
| Pereira et al., 2018 | 1.00 | 1.0000 | 1 | 1.0% | 1.00 [-0.96; 2.96] |
| Pereira et al., 2018 | 1.00 | 1.0000 | 1 | 1.0% | 1.00 [-0.96; 2.96] |
| Pereira et al., 2018 | 5.00 | 2.2361 | 1 | 0.4% | 5.00 [0.62; 9.38] |
| Santos-Silva et al., 2006 | 4.00 | 2.0000 | 1 | 0.5% | 4.00 [0.08; 7.92] |
| Santos-Silva et al., 2006 | 1.00 | 1.0000 | 1 | 1.0% | 1.00 [-0.96; 2.96] |
| Santos-Silva et al., 2006 | 1.00 | 1.0000 | 1 | 1.0% | 1.00 [-0.96; 2.96] |
| Santos-Silva et al., 2006 | 7.00 | 2.6458 | 1 | 0.3% | 7.00 [1.81; 12.19] |
| Santos-Silva et al., 2006 | 4.00 | 2.0000 | 1 | 0.5% | 4.00 [0.08; 7.92] |
| Santos-Silva et al., 2006 | 3.00 | 1.7321 | 1 | 0.6% | 3.00 [-0.39; 6.39] |
| Norte et al., 2012 | 1.00 | 1.0000 | 1 | 1.0% | 1.00 [-0.96; 2.96] |
| Palomar et al., 2016 | 7.00 | 2.6458 | 1 | 0.3% | 7.00 [1.81; 12.19] |
| Palomar et al., 2016 | 1.00 | 1.0000 | 1 | 1.0% | 1.00 [-0.96; 2.96] |
| Palomar et al., 2016 | 2.00 | 1.4142 | 1 | 0.7% | 2.00 [-0.77; 4.77] |
| Palomar et al., 2016 | 4.00 | 2.0000 | 1 | 0.5% | 4.00 [0.08; 7.92] |
| Palomar et al., 2016 | 1.00 | 1.0000 | 1 | 1.0% | 1.00 [-0.96; 2.96] |
| Palomar et al., 2016 | 1.00 | 1.0000 | 1 | 1.0% | 1.00 [-0.96; 2.96] |
| Palomar et al., 2016 | 8.00 | 2.8284 | 1 | 0.3% | 8.00 [2.46; 13.54] |
| Palomar et al., 2016 | 2.00 | 1.4142 | 1 | 0.7% | 2.00 [-0.77; 4.77] |
| Palomar et al., 2016 | 2.00 | 1.4142 | 1 | 0.7% | 2.00 [-0.77; 4.77] |
| Palomar et al., 2016 | 1.00 | 1.0000 | 1 | 1.0% | 1.00 [-0.96; 2.96] |
| Palomar et al., 2015 | 1.00 | 1.0000 | 1 | 1.0% | 1.00 [-0.96; 2.96] |
| England et al., 2016 | 0.10 | 0.3162 | 10 | 1.5% | 0.10 [-0.10; 0.30] |
| Palomar et al., 2016 | 2.82 | 1.6787 | 11 | 1.3% | 2.82 [1.83; 3.81] |
| England et al., 2016 | 0.42 | 0.6455 | 12 | 1.4% | 0.42 [0.05; 0.78] |
| Palomar et al., 2017 | 0.28 | 0.5301 | 121 | 1.5% | 0.28 [0.19; 0.38] |
| Pereira et al., 2018 | 1.50 | 1.2247 | 2 | 1.1% | 1.50 [-0.20; 3.20] |
| Pereira et al., 2018 | 0.50 | 0.7071 | 2 | 1.3% | 0.50 [-0.48; 1.48] |
| Pereira et al., 2018 | 3.50 | 1.8708 | 2 | 0.8% | 3.50 [0.91; 6.09] |
| Santos-Silva et al., 2006 | 1.50 | 1.2247 | 2 | 1.1% | 1.50 [-0.20; 3.20] |
| Santos-Silva et al., 2006 | 9.50 | 3.0822 | 2 | 0.4% | 9.50 [5.23; 13.77] |
| Santos-Silva et al., 2006 | 7.50 | 2.7386 | 2 | 0.5% | 7.50 [3.70; 11.30] |
| Santos-Silva et al., 2006 | 1.50 | 1.2247 | 2 | 1.1% | 1.50 [-0.20; 3.20] |
| Santos-Silva et al., 2006 | 1.00 | 1.0000 | 2 | 1.2% | 1.00 [-0.39; 2.39] |
| Santos-Silva et al., 2006 | 3.00 | 1.7321 | 2 | 0.8% | 3.00 [0.60; 5.40] |
| Palomar et al., 2016 | 1.00 | 1.0000 | 2 | 1.2% | 1.00 [-0.39; 2.39] |
| Palomar et al., 2016 | 1.50 | 1.2247 | 2 | 1.1% | 1.50 [-0.20; 3.20] |
| Palomar et al., 2016 | 3.00 | 1.7321 | 2 | 0.8% | 3.00 [0.60; 5.40] |
| Palomar et al., 2016 | 1.00 | 1.0000 | 2 | 1.2% | 1.00 [-0.39; 2.39] |
| Palomar et al., 2015 | 0.50 | 0.7071 | 2 | 1.3% | 0.50 [-0.48; 1.48] |
| Palomar et al., 2015 | 0.62 | 0.7906 | 24 | 1.5% | 0.62 [0.31; 0.94] |
| Santos-Silva et al., 2006 | 0.67 | 0.8165 | 3 | 1.3% | 0.67 [-0.26; 1.59] |
| Palomar et al., 2016 | 6.00 | 2.4495 | 3 | 0.7% | 6.00 [3.23; 8.77] |
| Palomar et al., 2015 | 1.00 | 1.0000 | 3 | 1.3% | 1.00 [-0.13; 2.13] |
| Palomar et al., 2016 | 2.50 | 1.5811 | 4 | 1.1% | 2.50 [0.95; 4.05] |
| Palomar et al., 2015 | 1.50 | 1.2247 | 4 | 1.2% | 1.50 [0.30; 2.70] |
| Palomar et al., 2015 | 1.25 | 1.1180 | 4 | 1.3% | 1.25 [0.15; 2.35] |
| Pereira et al., 2018 | 0.02 | 0.1429 | 49 | 1.5% | 0.02 [-0.02; 0.06] |
| Santos-Silva et al., 2006 | 4.00 | 2.0000 | 5 | 1.0% | 4.00 [2.25; 5.75] |
| Palomar et al., 2016 | 2.00 | 1.4142 | 5 | 1.2% | 2.00 [0.76; 3.24] |
| Palomar et al., 2016 | 1.60 | 1.2649 | 5 | 1.3% | 1.60 [0.49; 2.71] |
| Palomar et al., 2016 | 1.20 | 1.0954 | 5 | 1.3% | 1.20 [0.24; 2.16] |
| Santos-Silva et al., 2006 | 4.67 | 2.1602 | 6 | 1.0% | 4.67 [2.94; 6.40] |
| Palomar et al., 2015 | 0.17 | 0.4082 | 6 | 1.4% | 0.17 [-0.16; 0.49] |
| Pereira et al., 2018 | 2.71 | 1.6475 | 7 | 1.2% | 2.71 [1.49; 3.93] |
| Hoffman et al., 2020 | 0.05 | 0.2122 | 7354 | 1.5% | 0.05 [0.04; 0.05] |
| Foley-Fisher et al., 2012 | 0.25 | 0.5000 | 8 | 1.4% | 0.25 [-0.10; 0.60] |
| England et al., 2016 | 0.62 | 0.7906 | 8 | 1.4% | 0.62 [0.08; 1.17] |
| England et al., 2016 | 0.12 | 0.3536 | 8 | 1.5% | 0.12 [-0.12; 0.37] |
| Palomar et al., 2015 | 0.25 | 0.5000 | 8 | 1.4% | 0.25 [-0.10; 0.60] |
| Norte et al., 2012 | 0.24 | 0.4942 | 86 | 1.5% | 0.24 [0.14; 0.35] |
| Total (95% CI) | | | 7811 | 64.2% | 1.10 [0.88; 1.33] |
| Heterogeneity: $Tau^2 = 0.3425$; $Chi^2 = 354.63$, df = 62 ($P < 0.01$); $I^2 = 83\%$ | | | | | |
| **Landscape = Continental** | | | | | |
| Kolodziejek et al., 2014 | 1.00 | 1.0000 | 1 | 1.0% | 1.00 [-0.96; 2.96] |
| Capek et al., 2014 | 0.01 | 0.1109 | 325 | 1.5% | 0.01 [0.00; 0.02] |
| Capek et al., 2014 | 0.03 | 0.1733 | 333 | 1.5% | 0.03 [0.01; 0.05] |
| Sandor et al., 2017 | 0.02 | 0.1400 | 51 | 1.5% | 0.02 [-0.02; 0.06] |
| Hornok et al., 2013 | 0.00 | 0.0639 | 734 | 1.5% | 0.00 [-0.00; 0.01] |
| Capek et al., 2014 | 0.11 | 0.3273 | 84 | 1.5% | 0.11 [0.04; 0.18] |
| Capek et al., 2014 | 0.01 | 0.1054 | 90 | 1.5% | 0.01 [-0.01; 0.03] |
| Total (95% CI) | | | 1618 | 9.8% | 0.07 [-0.39; 0.53] |
| Heterogeneity: $Tau^2 = 0.3425$; $Chi^2 = 17.55$, df = 6 ($P < 0.01$); $I^2 = 66\%$ | | | | | |
| **Landscape = Steppic** | | | | | |
| Palomar et al., 2016 | 24.00 | 4.8990 | 1 | 0.1% | 24.00 [14.40; 33.60] |
| Dantas-Torres et al., 2011 | 9.71 | 3.1154 | 17 | 1.1% | 9.71 [8.22; 11.19] |
| Total (95% CI) | | | 18 | 1.2% | 10.22 [8.38; 12.06] |
| Heterogeneity: $Tau^2 = 0.3425$; $Chi^2 = 8.32$, df = 1 ($P < 0.01$); $I^2 = 88\%$ | | | | | |
| **Landscape = Anatolian** | | | | | |
| Keskin and Erciyas-Yavuz, 2019 | 0.00 | 0.0446 | 1004 | 1.5% | 0.00 [-0.00; 0.00] |
| Keskin and Erciyas-Yavuz, 2019 | 0.15 | 0.3862 | 114 | 1.5% | 0.15 [0.08; 0.22] |
| Keskin and Erciyas-Yavuz, 2019 | 0.01 | 0.0756 | 175 | 1.5% | 0.01 [-0.01; 0.02] |
| Keskin and Erciyas-Yavuz, 2019 | 0.21 | 0.4588 | 19 | 1.5% | 0.21 [0.00; 0.42] |
| Keskin and Erciyas-Yavuz, 2019 | 0.02 | 0.1351 | 274 | 1.5% | 0.02 [0.00; 0.03] |
| Keskin and Erciyas-Yavuz, 2019 | 0.00 | 0.0579 | 298 | 1.5% | 0.00 [-0.00; 0.01] |
| Keskin and Erciyas-Yavuz, 2019 | 0.58 | 0.7620 | 31 | 1.5% | 0.58 [0.31; 0.85] |
| Keskin and Erciyas-Yavuz, 2019 | 0.06 | 0.2425 | 34 | 1.5% | 0.06 [-0.02; 0.14] |
| Keskin and Erciyas-Yavuz, 2019 | 0.00 | 0.0640 | 489 | 1.5% | 0.00 [-0.00; 0.01] |
| Keskin and Erciyas-Yavuz, 2019 | 0.01 | 0.1204 | 69 | 1.5% | 0.01 [-0.01; 0.04] |
| Keskin and Erciyas-Yavuz, 2019 | 0.03 | 0.1622 | 76 | 1.5% | 0.03 [-0.01; 0.06] |
| Keskin and Erciyas-Yavuz, 2019 | 0.01 | 0.0762 | 860 | 1.5% | 0.01 [0.00; 0.01] |
| Keskin and Erciyas-Yavuz, 2019 | 0.44 | 0.6667 | 9 | 1.4% | 0.44 [0.01; 0.88] |
| Keskin and Erciyas-Yavuz, 2019 | 0.00 | 0.0471 | 903 | 1.5% | 0.00 [-0.00; 0.01] |
| Total (95% CI) | | | 4355 | 20.5% | 0.10 [-0.21; 0.41] |
| Heterogeneity: $Tau^2 = 0.3425$; $Chi^2 = 51.66$, df = 13 ($P < 0.01$); $I^2 = 75\%$ | | | | | |
| **Landscape = Boreal** | | | | | |
| Movila et al., 2013 | 0.18 | 0.4264 | 11 | 1.5% | 0.18 [-0.07; 0.43] |
| Movila et al., 2013 | 0.06 | 0.2425 | 17 | 1.5% | 0.06 [-0.06; 0.17] |
| Movila et al., 2013 | 0.67 | 0.8165 | 3 | 1.3% | 0.67 [-0.26; 1.59] |
| Total (95% CI) | | | 31 | 4.2% | 0.25 [-0.47; 0.97] |
| Heterogeneity: $Tau^2 = 0.3425$; $Chi^2 = 2.29$, df = 2 ($P = 0.32$); $I^2 = 12\%$ | | | | | |
| **Total (95% CI)** | | | **13833** | **100.0%** | **1.18 [0.85; 1.51]** |

Heterogeneity: $Tau^2 = 1.9244$; $Chi^2 = 897.66$, df = 88 ($P < 0.01$); $I^2 = 90\%$
Residual heterogeneity: $Tau^2 = 0.3425$; $Chi^2 = 434.44$, df = 84 ($P < 0.01$); $I^2 = 81\%$
Test for subgroup differences: $Chi^2 = 138.23$, df = 4 ($P < 0.01$)

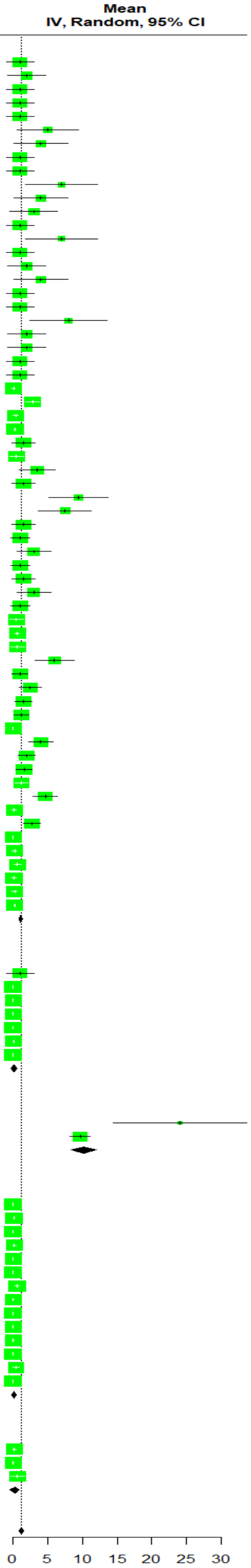


***Figure S7**. subgroup meta analysis of immatures parasitic load using landscape as the grouping variable*
